# Apicomplexan F-actin is required for efficient nuclear entry during host cell invasion

**DOI:** 10.1101/646463

**Authors:** Mario Del Rosario, Javier Periz, Georgios Pavlou, Oliver Lyth, Fernanda Latorre-Barragan, Sujaan Das, Gurman S. Pall, Johannes Felix Stortz, Leandro Lemgruber, Jake Baum, Isabelle Tardieux, Markus Meissner

**Author notes:** **Abbreviation: CD** Cytochalasin-D, **CLEM** corelative light and electron microscopy, **TJ** tight junctional ring, **SR-SIM** Super Resolution – Structure Illumination Microscopy, **Cb** Chromobody, **FP** Fluorescent protein.

## Abstract

The obligate intracellular parasites *Toxoplasma gondii* and *Plasmodium spp*. invade host cells by injecting a protein complex into the membrane of the targeted cell that bridges the two cells through the assembly of a ring-like junction. This circular junction stretches while the parasites applies a traction force to pass through; a step that typically concurs with transient constriction of the parasite body. Here we show that the junction can oppose resistance to the passage of the parasite’s nucleus. Super-resolution microscopy and real time imaging highlighted an F-actin pool at the apex of pre-invading parasite, an F-actin ring at the junction area during invasion but also networks of perinuclear and posteriorly localized F-actin. Mutant parasites with dysfunctional acto-myosin showed significant decrease of junctional and perinuclear F-actin and are coincidently affected in nuclear passage through the junction. We propose that the F-actin machinery eases nuclear passage by stabilising the junction and pushing the nucleus through the constriction, providing first evidence for a dual contribution of actin-forces during host cell invasion by apicomplexan parasites.

## Introduction

The phylum Apicomplexa consists of more than 5000 species, most of them obligate intracellular parasites, including important human and veterinary pathogens, such as *Plasmodium* (malaria) or *Toxoplasma* (toxoplasmosis).

During their complex life cycles, apicomplexan parasites move through different environments to disseminate within and between hosts and to invade their host cell (Meissner, Ferguson et al., 2013). Therefore, the invasive stages, called zoites evolved a unique invasion device, consisting of unique secretory organelles and the parasites acto-myosin system, the glideosome, localised in the narrow space (∼30nm) between the plasma membrane and the Inner Membrane Complex (IMC) (Frenal, Dubremetz et al., 2017). Zoites actively enter the host cell by establishing a tight junctional ring (TJ) at the point of contact between the two cells. The TJ is assembled by the sequential secretion of unique secretory organelles (micronemes and rhoptries), leading to the insertion of rhoptry neck proteins (RONs) into the host cell Plasma Membrane (PM) and underneath (Besteiro, Dubremetz et al., 2011). On the extracellular side, the exposed domain of the RON2 member binds the micronemal transmembrane protein AMA1 exposed on the parasite surface, resulting in the formation of a stable, junctional complex (Besteiro et al., 2011). The TJ is further anchored to the host cell cortex by *de novo* formation of F-actin through the recruitment of actin-nucleating proteins (Bichet, Joly et al., 2014), (Guerin, Corrales et al., 2017). During host cell invasion the parasites use their acto-myosin motor to pass through the TJ. However, the exact role and orientation of the parasite’s acto-myosin system is still under debate (Tardieux & Baum, 2016) and intriguingly, mutants for key component of this system show residual motile and invasive capacities (Egarter, Andenmatten et al., 2014, Gras, Jackson et al., 2017, Whitelaw, Latorre-Barragan et al., 2017), the latter reflecting in large part an alternative and host-cell actin-dependant mode of entry (Bichet, Touquet et al., 2016). According to the glideosome model the force generated for motility and invasion relies exclusively on F-actin polymerised at the apical tip of the parasite by the action of Formin-1 and translocated within the narrow space (∼30 nm) between the Inner Membrane Complex (IMC) and PM of the parasite (Tosetti, Dos Santos Pacheco et al., 2019). In support of this model, was the detection of parasite F-actin underneath the junction formed by invading parasites when using an antibody preferentially recognising apicomplexan F-actin. Furthermore, the detection of cytosolic locations, predominantly around the nucleus (Angrisano, Riglar et al., 2012) suggest additional roles of this cytoskeletal protein during invasion.

While it was assumed a major role of F-actin in driving Apicomplexa zoite gliding motility and cell invasion, recent studies demonstrated the pivotal role of F-actin in multiple other processes, such as apicoplast inheritance (Andenmatten, Egarter et al., 2013), dense granule motility (Heaslip, Nelson et al., 2016) and likely nuclear functions through the control of expression of virulence genes in malaria parasites (Zhang, Huang et al., 2011). However, building a comprehensive model for F-actin dynamics, localisation and function in apicomplexan parasite has been hampered for decades by the lack of tools enabling reliable F-actin detection. While some studies suggested that F-actin is interacting with subpellicular microtubules (Patron, Mondragon et al., 2005) and/or the subpellicular matrix of the IMC (Hliscs, Millet et al., 2015) (Yasuda, Yagita et al., 1988), they were seen with some scepticism, since they were incompatible with the prevailing model for the glideosome.

With the adaptation of nanobodies specifically recognising F-actin, this limitation has been overcome (Periz, Whitelaw et al., 2017) leading to the identification of distinct cytosolic networks of dynamic actin in both *T. gondii* and *P. falciparum* (Periz et al., 2017) (Tosetti et al., 2019)(Stortz et al., BioRXiv2018). Recently, three polymerisation centres have been described, correlating with the location of Formins-1,2 and 3 in *T. gondii* (Tosetti et al., 2019). Formins-2 and -3 appear to have overlapping functions for material exchange and formation of an intravacuolar network in intracellular parasites; Formin-1 (FH-1) has been explicitly implicated in the apical polymerisation of F-actin required for gliding motility and cell invasion(Tosetti et al., 2019). In this study, although not directly shown, the authors suggested that FH-1 polymerises F-actin at the apical tip of the parasite that is subsequently transported by the action of the glideosome within the sub-alveolar space to the posterior pole.

Using well standardised transgenic parasites expressing Cb, we previously identified two F-actin polymerisation centres from where most of the F-actin flow occurs in intracellular parasites (Periz et al., 2017). One polymerisation centre can be found at the apical tip, corresponding well to the described location of Formin-1, and one close to the parasite’s Golgi, corresponding to the location of Formin-2 (Tosetti et al., 2019)(Stortz et al., BioRXiv2018). Interestingly, at least in intracellular parasites, the majority of F-actin dynamics appears to occur within the cytosol of the parasite close to the parasite’s Golgi, forming a continuous dynamic flow from centre to the periphery of the parasite and, from the apical to the basal pole(Periz et al., 2017).

In good agreement with a cytosolic oriented F-actin system, previous reports demonstrated parasite F-actin to be associated with the sub-pellicular microtubules that are connected to and stabilise the IMC (Patron et al., 2005) (Yasuda et al., 1988) (Shaw, Compton et al., 2000) (Hliscs et al., 2015). Furthermore, gliding associated proteins identified via co-immunoprecipitation with the glideosome, were recently found to play important roles in stabilising the IMC and directly connecting it to the subpellicular microtubules (Harding, Egarter et al., 2016) (Harding, Gow et al., 2019), suggesting that a close connection exists between parasite F-actin, the glideosome and the subpellicular network.

Here we compared F-actin dynamics in WT and mutant parasites for the acto-myosin system and correlated its dynamic location with the phenotypic consequences during host cell invasion. We identified the nucleus as a major obstacle for efficient host cell invasion that needs to be squeezed through the junction in a F-actin dynamic manner.

Our analysis strongly suggests a push-and-pull mechanism for nuclear entry during host cell invasion, in analogy to the dynamics observed during migration of other eukaryotes through a constricted environment (McGregor, Hsia et al., 2016).

## Results

### F-actin flow is mainly organised within the cytosol of extracellular parasites and is modulated by Calcium and cGMP signalling

We previously imaged F-actin flow in intracellular parasites and were able to discriminate 2 major polymerisation centres, one at the apical tip and one close to the apicoplast, where F-actin is formed and subsequently transported to the posterior end of the parasite (Periz et al., 2017). Indeed, recent studies on *Toxoplasma* and *Plasmodium* demonstrated that Formin-2 is localised to the apicoplast of the parasite and required for most of the intracellular F-actin dynamic (Stortz et al., BioRXiv2019; (Tosetti et al., 2019)), while Formin-1 is localised at the apical tip, where it is thought to be exclusively required for parasite motility and invasion. In good agreement, in intracellular parasites F-actin appears to be formed at two polymerisation centres, localised at the apical tip and close to the Golgi region of the parasite, indicating that FH-1 and FH-2 are acting as nucleators during intracellular parasite development. Intriguingly, the majority of F-actin dynamics in intracellular parasites occurs within the cytosol of the parasite. (Periz et al., 2017). We were interested to determine if during the transient extracellular life and at the time of cell invasion, the patterns of actin dynamics differs from the intracellular ones and how they were impacted upon disruption of the glideosome or factors critically involved in F-actin regulation.

Since transient transfections is typically associated with overexpression of the target protein hence with significant uncontrolled impact on a dynamic equilibrium (Periz et al., 2017), we generated *T. gondii* lines stably expressing Cb-EmeraldFP at comparable levels and ensured that its expression does not lead to phenotypic alterations and misleading analysis. We established transgenic parasites expressing Cb-EmeraldFP in WT parasites, a null mutant for myosin A (MyoA), the core motor of the glideosome(Andenmatten et al., 2013) and a conditional mutant for the critical actin regulator, actin depolymerisation factor ADF, *adf*cKD (Mehta & Sibley, 2011). Using live 3D-structure illumination microscopy (3D-SIM), we compared F-actin dynamics and found two discernible F-actin polymerisation centres (Fig.1A) as previously seen in intracellular parasites (Periz et al., 2017). Time lapse analysis demonstrated that the majority of F-actin dynamics occurs in the cytosol of the parasite with some F-actin flow detectable at the periphery. In good agreement with the localisation of FH-1 and FH-2, two polymerisation centres can be detected. F-actin flow starts from the apical tip (1^st^ polymerisation centre, yellow arrow in Fig. 1A) to the second polymerisation centre close to the Golgi (red arrow, Fig.1A) to the posterior pole of the parasite (Fig.1A, Movie S1). While disruption of MyoA did not result in significant changes of F-actin localisation or dynamics in resting parasites (Fig.1A, MovieS1), depletion of ADF completely abrogated actin dynamics, F-actin accumulation being observed at both apical and posterior parasite poles (Fig.1A, MovieS1).

**Figure 1.**
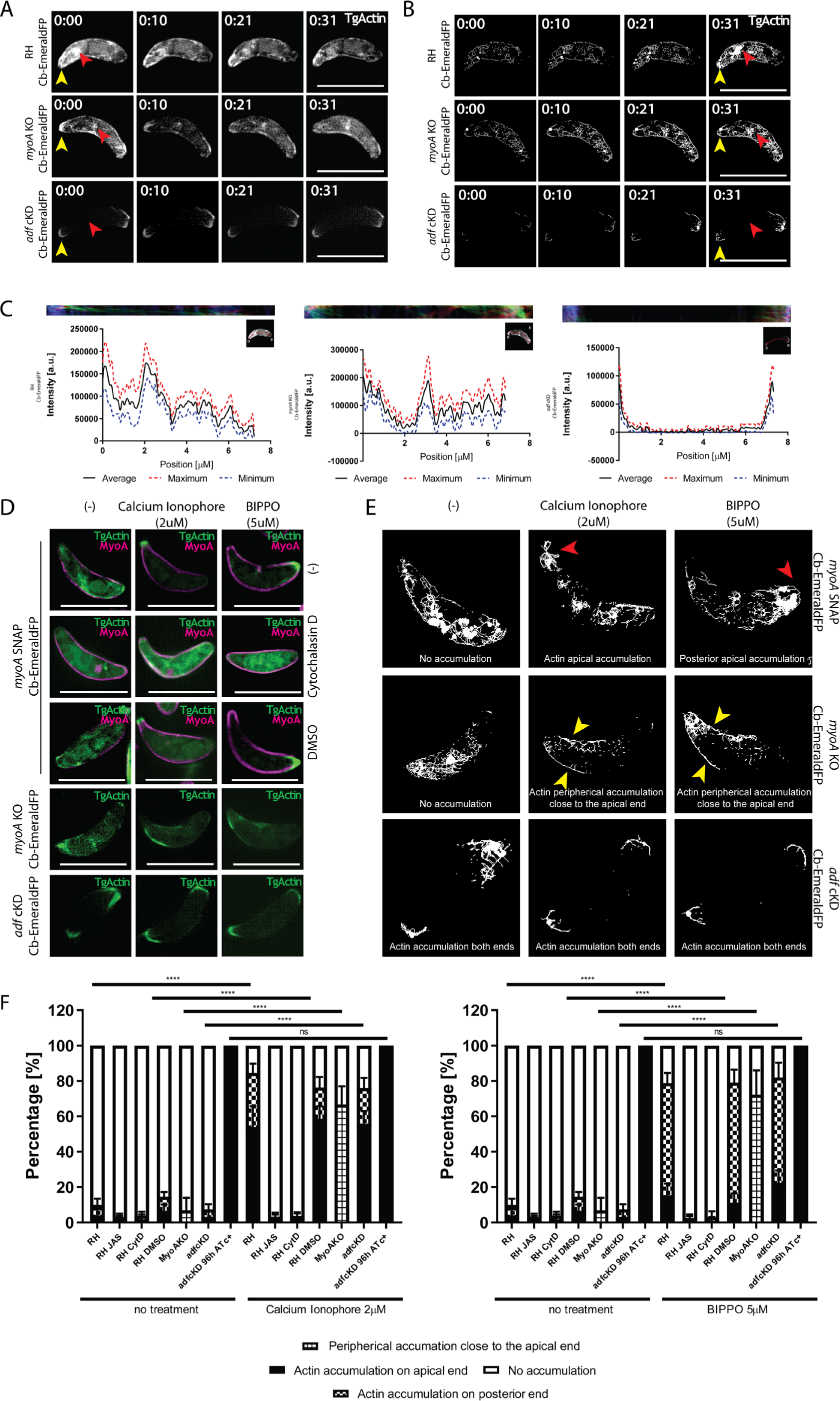
F-actin, formed at two major nucleation centres forms a dynamic, continuous network within the cytosol of the parasite. **A.** Stills depicting actin flow (see movie S1) in extracellular RH, *myoA* KO and *adf* KD parasites expressing Cb EmeraldFP. Continuous F-actin flow can be seen from the apical tip, to the Golgi region towards the posterior pole of the parasite. No difference can be observed between RH and *myoA* KO. For *adf* KD, the flow is completely abrogated. Red arrowhead marks the Golgi region, while the yellow arrowhead marks the apical nucleation centre. **B.** Skeletonisation processing of **(A)** depicting areas where F-actin dynamics/flow is prevalent. Individual signals for F-actin form a continuous, dynamic network that connects apical and posterior pole of the parasite. Red arrowhead marks the Golgi region nucleation centre, while the yellow arrowhead marks the apical nucleation centre. **C.** Kymograph analysis of **(A)** and **(B)**. The kymograph was generated by tracing a line in the middle of the parasite’s body (red line). The colour-coded kymograph represents forward movement (red), backwards movement (green) and static F-actin (blue). The results demonstrate continuous exchange of F-actin from the apical to the posterior pole of the parasite within the cytosol of the parasite. This cytosolic exchange is similar in RH and *myoA* KO parasites, while completely abolished in the case of *adf* KD. **D.** Top three rows: Parasites expressing Cb EmeraldFP along with MyoA-SNAP before and after Ca^2+^-Ionophore (A23187) or BIPPO treatment. The parasites were also treated with 2 µM of Cytochalasin D or DMSO (as control) for 30 min. After addition of A23187, preferential relocalisation of actin can be observed at the apical tip, while addition of BIPPO caused F-actin accumulation at the basal end. 4th row: *myoA*KO parasites expressing Cb-EmeraldFP before and after treatment with A23187 and BIPPO. While no apical or basal accumulation of F-actin is observed, some peripheral location of F-actin occurs in presence of A23187 or BIPPO, preferentially in the apical half of the parasite. Bottom row: *adf* cKD parasites expressing Cb-EmeraldFP before and after treatment with Ca^2+^ Ionophore or BIPPO. No apparent change in F-actin localisation can be seen upon treatment. Representative still from Movie S2. **E.** Skeletonization of movies shown in **D**. Before addition of A23187 or BIPPO, RH and *myoA* KO parasites behave similar with no accumulation of F-actin at the apical tip. After treatment with A23187, preferential accumulation of F-actin at the apical tip can be observed for RH, while BIPPO presented preferential accumulation in the basal end. In *myoA* KO, some relocalisation of actin is observed at the periphery of the parasite. In the case of *adf* cKD, no relocalisation occurs. **F.** Quantification of actin accumulation before and after treatment with Ca^2+^ Ionophore or BIPPO. The parasites were counted for F-actin accumulation after adding Ca^2+^ Ionophore or BIPPO. Numbers were generated by counting total number of parasites and then number of parasites with Actin accumulation on the apical or basal tip. A minimum of 200 parasites were counted upon 3 biological replicates. Two-way ANOVA was used for statistical analysis and Tukey’s multiple comparisons test. p value <0.0001. Numbers indicate minutes:seconds; and Scale bars represent 5 µm.

To map the actin structures visualised with the chromobody, we employed skeletonization processing(Arganda-Carreras, Fernandez-Gonzalez et al., 2010), which converts F-actin signals into individual pixels, that can be traced to obtain information about the localisation and dynamics of F-actin. The resulting skeletonised movie can be projected to show where the majority of (detectable) F-actin dynamics occur during the timeline of the movie (Fig. 1B, Movie S1). This analysis confirmed that parasite F-actin formed at the apical tip (yellow arrowhead) and Golgi region (red arrowhead), converge within the cytosol of the parasite and reach the basal pole of the parasite. Interestingly, both peripherical and cytoplasmic flow appear to be connected and do not occur independently from each other (Fig.1B). Note that both WT and *myoA*KO parasites displayed similar signals whereas the ADF-conditional mutant did not. Next, kymograph analysis was performed to analyse individual F-actin flow events using KymographClear and KymographDirect (Mangeol, Prevo et al., 2016) confirming the presence of a continuous dynamic F-actin network in cytosolic location that connects the apical pole, Golgi area and basal pole of the parasite, in good agreement with the location of the 3 Formins previously described in *T. gondii* (Tosetti et al., 2019). KymographClear is a FIJI plugin that generates a 3 colour-coded kymograph, where each colour labels either forward movement (red), backward movement (green) or static (blue) as shown (Fig. 1C and Fig. S1). This kymograph can then be read into the KymographDirect software, which uses automatic detection to trace trajectories of moving particles in the image. Values are then assigned to these trajectories based on both intensity and speed (Mangeol et al., 2016). The intensity can be assessed in specified tracks over time as the average intensity over time shown in Fig. 1C. Both the RH and *myoA* KO, presented dynamic signals that can be seen at the apical pole (close to the point 0), Golgi region (between 2 and 3 µM) and basal pole. In the case of the *adf* KD, only static signals could be detected at the apical and basal pole of the parasite, confirming the lack of F-actin dynamics in these mutants.

Next, we used this analysis to investigate intensity profiles of actin signal travelling in the periphery of the parasite (Figure S1 B-D). Interestingly, actin dynamics along the periphery is similar between WT and *myoA* KO with bi-directional actin flow events as revealed by the kymographs, while confirming previous results with the *adf* KD mutants, which shows a lack of F-actin dynamics across the entire parasites body (first half (-), Figure S1 A-C).

Together, these data point that as expected and similarly to intracellular parasites, the F-actin is highly dynamic in extracellular parasites displaying two major polymerisation centres at the apical tip and Golgi-region of the parasite respectively. Of note, this analysis underlines that there is actin flow coming from the cytoplasm towards the periphery and shaping a highly dynamic network that connects the apical and basal pole of the parasite, therefore independently of any F-actin network positioned between the zoite PM and IMC.

Tosetti et al. suggested that activation of a Ca^2+^-signalling cascade would lead to increased retrograde transport of F-actin, eventually accumulating posteriorly (Tosetti et al., 2019). However we found that treatment of parasites with the calcium-ionophore A23187 induced anterior accumulation of F-actin in wild type parasites in the majority of cases (Fig.1D,E,F; Fig. S1A; Movie S2), while BIPPO triggered a predominantly basal F-actin accumulation (Fig.1D, E,F; Fig. S1A; Movie S2). BIPPO is a drug capable of inhibiting 3′,5′-cyclic nucleotide phosphodiesterases (PDEs) responsible in blocking the breakdown of cyclic nucleotides such as cAMP and cGMP. This mechanism is believed to be responsible in the activation of microneme secretion and egress by modulating the signal pathway of PKG-dependant processes(Howard, Harvey et al., 2015).

Once MyoA null mutants were submitted to treatments with A23187 or BIPPO, the F-actin remained rather evenly distributed within the cytoplasm albeit with some peripheral enrichment in the first half of the parasite closer to the apical end (Fig.1D,E,F; Movie S2). In contrast, depletion of ADF led to the expected phenotype with F-actin accumulating at the apical and basal pole of the parasite, independently of A23187 or BIPPO addition (Fig.1D,E,F; Movie S2). Together these data suggest that calcium-signalling regulates F-actin dynamics, potentially by stimulating the activity of Formin-1 followed by F-actin transport to the posterior pole of the parasite by the action of Myosin A.

In conclusion, similar to intracellular parasites, extracellular parasites show impressive F-actin dynamics within the cytosol. In parasite mutants for ADF, this dynamic is abolished, and F-actin accumulates at the apical and basal pole, independent of triggering a Calcium-signalling cascade. In contrast, disruption of MyoA results in normal F-actin dynamics, but upon Calcium-signalling F-actin remains more diffusely localised within the cytosol of the parasite with some accumulation at the periphery.

### The acto -myosin system is required to ensure efficient invasion

To investigate whether the impressive cytoplasmic F actin dynamics could assist the parasite over the invasion of the target cell, namely when it fits through the junction and constrict (Fig.2A,B), we first measured the average diameter of parasites prior and during invasion. We observed a significant decrease at mid invasion, (prior: 2.5-3 μm and mid-invasion 1.5-2 μm) that attests significant body deformation and compression in order to fit through the junction (Fig.2A, B). In good agreement, analysis of the average diameter of the parasite’s nucleus, indicated that this organelle suffers a striking deformation, up to ∼50% of its normal diameter (i.e. less than 1 μm) while passing through the junction (Fig.2A, B). Since the acto-myosin system generates traction force at the junction to account for the invasive force, we investigated if interference causes changes, i.e. differences in the diameter of the TJ and/or stronger deformations of the parasite. While a slight reduction in the average TJ diameter was observed during invasion of *myoA* KO parasites (Fig.2C), the junctional ring appears remarkable constant upon interference with the acto-myosin system of the parasite.

**Figure 2.**
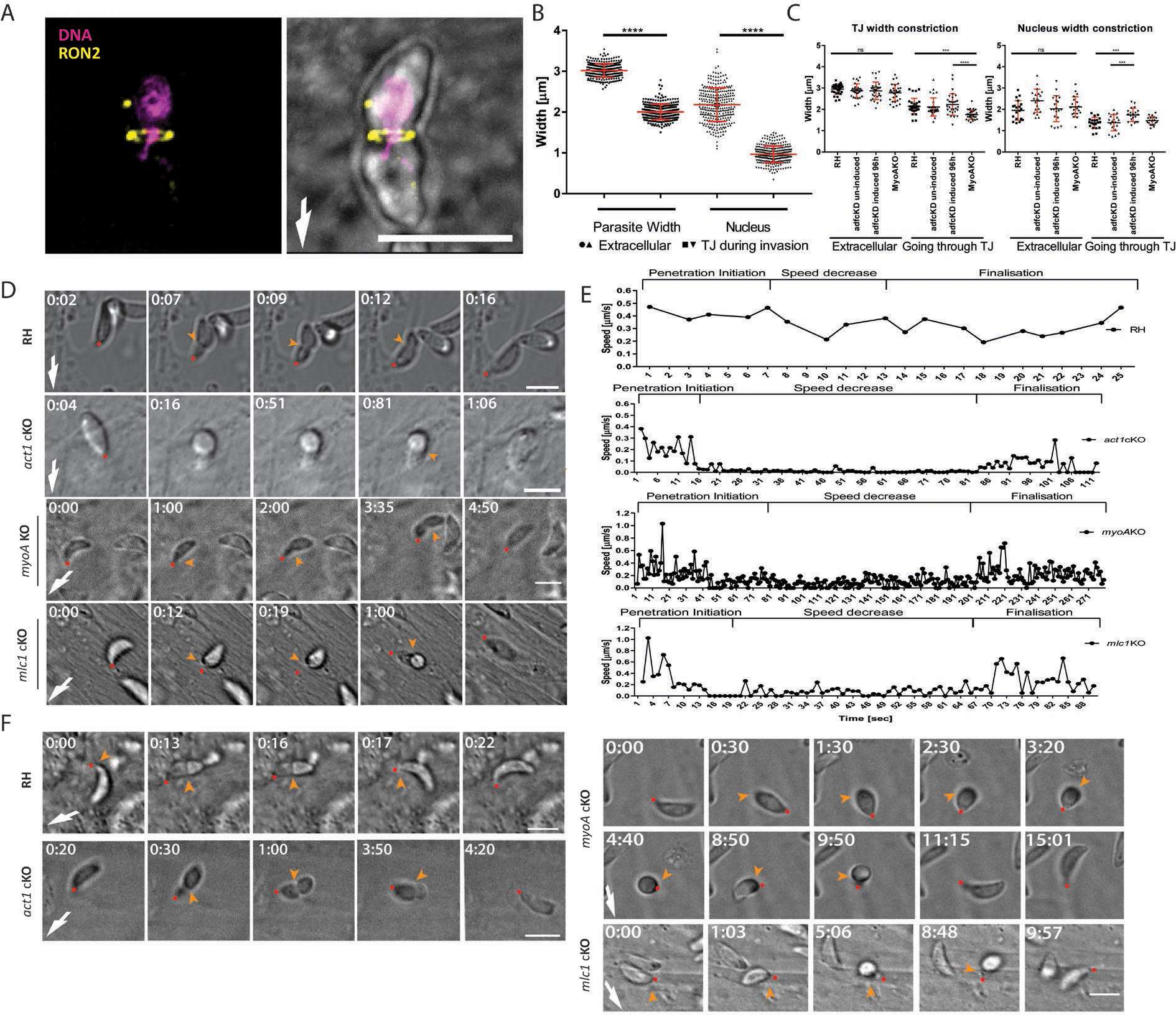
The nucleus is a limiting factor for host cell invasion. **A.** Wild Type RH parasites mid-invasion. Tight junction assay was done by allowing freshly egressed parasite to invade for 5 minutes, fixed with 4% PFA before labelling the TJ with anti-Ron2 (yellow) and DAPI (magenta) for nuclear staining. Scale bar represents 5 µm. **B.** Graph showing deformation of the whole parasite and its nucleus between invading and non-invading parasites. 100 parasites found mid-invasion were counted in triplicate and compared to extracellular, freshly egressed parasites. One-way ANOVA was used for statistical analysis. p value equals <0.0001. **C.** Graph showing deformation of the whole parasite and its nucleus between invading and non-invading parasites. Indicated parasites were analysed. For *adf* cKD induced and non-induced conditions were compared, as indicated. 30 parasites were counted for each condition in triplicate. One-way ANOVA was used for statistical analysis and Tukey’s multiple comparisons test. p value <0.0001. **D.** Time-lapse microscopy analysis of indicated parasites invading HFFs cells. Time was determined from the onset of invasion. **E.** Speed profiles of invading parasites shown in D. The relative displacement of the apical tip over time allowed speed calculations. Individual stages were named in regard to the invasion step (Penetration initiation, Speed decrease and finalisation). Invasion was analysed with Icy Image Processing Software (Pasteur Institut) using the Manual Tracking plugin by analysing each movie 10 times and averaging the measurements (for additional speed profiles see Fig.S2). **F.** Time-lapse analysis demonstrating abortive invasion of indicated parasites. Time was determined from the onset of invasion. White arrow points at invasion direction; yellow arrow points at tight junction; red dot follows the apical pole of the parasite; numbers indicate minutes:seconds; and Scale bars represent 5 µm.

Previously it was demonstrated that mutants for the acto-myosin system enter the host cell in a stop-and-go manner, leading to a prolonged invasion process with different features (Egarter et al., 2014, Mehta & Sibley, 2011, Whitelaw et al., 2017). While mutant parasites could be driven through the TJ by the host cell membrane outward projections in an uncompleted micropinocytosis-like process (Bichet et al., 2016), they could in some cases remain trapped in the TJ and die, probably due to compressive forces exerted on the embedded part of the parasite by the host cell membrane protrusions(Bichet et al., 2016). We infer from these observations that the tachyzoite likely needs to counteract compressive forces at the TJ to secure entry of its bulky organelles, such as the nucleus and hypothesised that the parasite acto-myosin is critically involved in nuclear entry.

We compared the invasion process of WT and mutant parasites in detail using time lapse analysis (Fig.2D, E, S2, Movie S3,4). Interestingly, the invasion process of WT parasites proceeded with an initial rapid invasion step, followed by a short pause when ∼1/3 of the body was inside the host cell, and another acceleration until the parasite was fully inside the host cell (Fig.2D,E, Movie S3). In contrast, whilst initial invasion steps appeared to be comparable between WT parasites and mutants for the acto-myosin system (*act1*cKO, *mlc1*cKO, *myoA*KO), an extended period then followed, where parasites stalled (up to 8 minutes), before invasion continued (Fig.2D,E, S2, MovieS4). In some cases, we also observed abortive invasion events, where the invading parasites terminally stalled before being pushed out from the TJ (Fig.2F, Movie S5). Interestingly, this stalling occurred approximately when the nucleus reached the TJ suggesting that in addition to providing the traction force that promotes entry through the TJ (Frenal et al., 2017), F-actin also assists the passage of the nucleus (and other bulky organelles, such as apical secretory complex) through the TJ, akin to other eukaryotes(Thiam, Vargas et al., 2016).

### F-actin accumulates at the posterior pole and in most cases at the junction

To correlate F-actin dynamics with the invasive behaviour of parasites, we expressed F-actin chromobodies in fusion with Emerald FP in *T. gondii* (Periz et al., 2017) WT and mutant parasites and compared the localisation and dynamics of F-actin when they invaded the host cells. To ensure specificity of Cb EmeraldFP for F-actin during the invasion process, we used a previously validated antibody raised against *P. falciparum* actin that preferentially recognises F-actin (Angrisano et al., 2012) and compared the staining obtained with this antibody on WT parasites with the staining obtained with Cb Emerald-expressing parasites. In both cases F-actin accumulation at the junction and the posterior pole was apparent (FigS3). Interestingly, while posterior accumulation of F-actin could always be detected, it was not systematic for the F-actin localized at the junction site (Fig.3A,E,F). We also ensured that expression of Cb Emerald did not cause any significant differences in the behaviour of parasites during invasion (Figure S3, see also (Periz et al., 2017)). Combining live cell and super-resolution microscopy, we were able to identify two situations in WT parasites: in the majority of cases (76%) we detected F-actin forming a ring coinciding with the TJ, while in ∼24% of all cases no significant accumulation was detectable, as confirmed by intensity plots (Fig.3A, B, C, S4, Movie S6). Importantly, as expected genetic interference with actin dynamics led to constrictions/TJ devoid of F-actin at the junction, as seen in the case of *adf* cKD (Fig.3A, B, Movie S6), demonstrating that F-actin accumulation at the junction requires correctly regulated F-actin dynamics, as suggested previously (Mehta & Sibley, 2011). Previously it was shown that WT parasites can enter the host cell through static or capped TJ depending on whether the TJ is respectively firmly or loosely anchored (Bichet et al., 2014). Analysis of parasites entering the host cell via capping shows F-actin at the TJ (Fig.3A, B, S4, Movie S6), confirming the finding that both invasion scenarios rely on the parasite F-actin-based traction force (Bichet et al., 2014).

**Figure 3.**
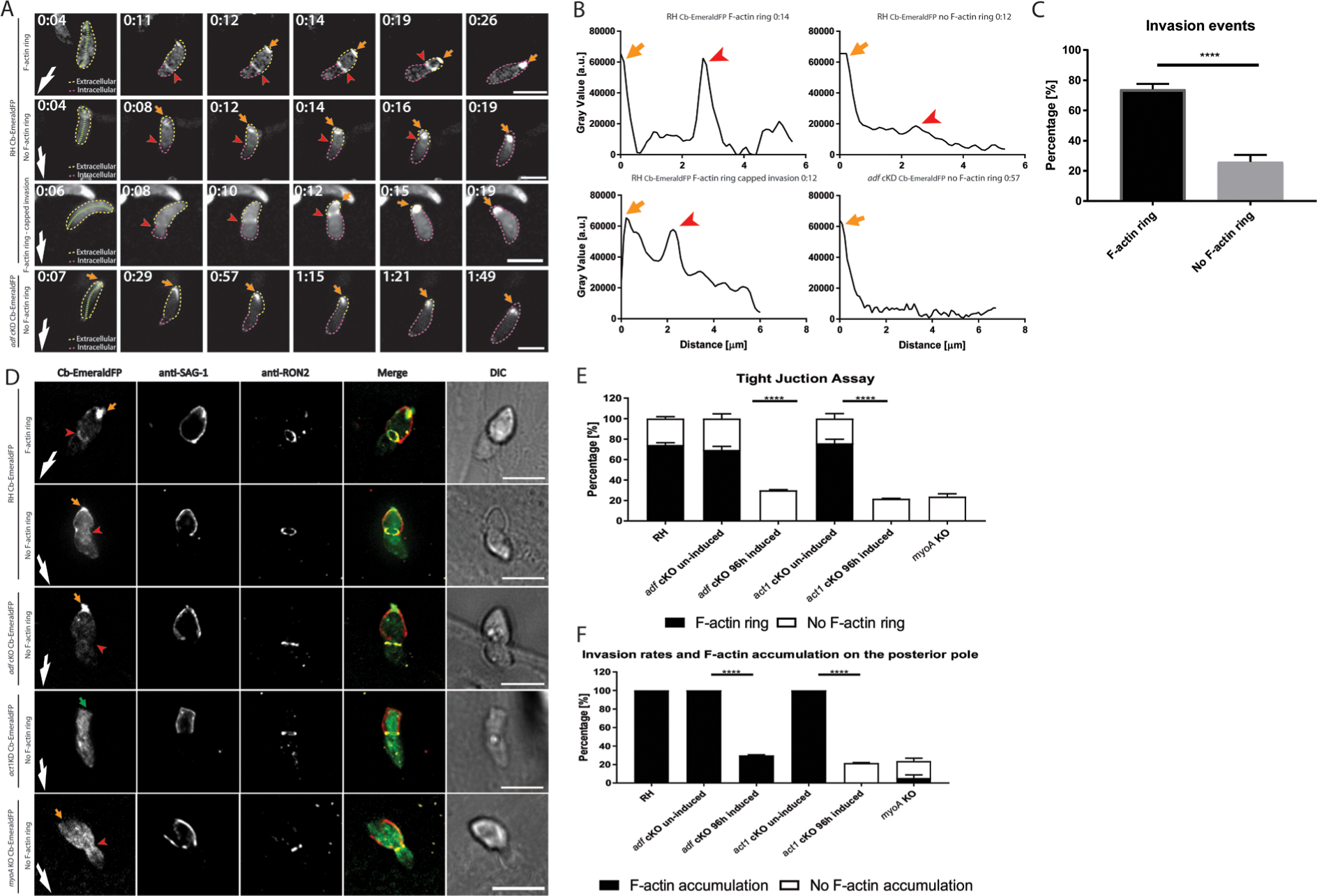
F-actin dynamics during *Toxoplasma gondii* invasion during fixed and live imaging. **A.** Time-lapse analysis depicting invading RH Cb-EmeraldFP parasites. During invasion, an F-actin ring can be observed at the TJ in the majority of cases (red arrowheads, first panel). Invasion can also occur without significant formation of an F-actin ring (second panel). During events of capped invasion, an F-actin ring is still formed (third panel). Invasion of *adf* cKD parasites occurs without formation of an F-actin ring despite clear deformation of the parasite body (fourth panel). During the entire invasion event F-actin accumulation is observed at the posterior pole of the parasite in all cases (orange arrows). Green segmented lines depict the area that was measured for generation of intensity plot profiles shown in **(B).** Yellow and purple dotted lines indicate the extra- and intracellular part of the parasite during the entry process respectively. **B.** Intensity plot profiles of the time lapse analysis shown in **(A).** Depicted plots correlate to half-invaded parasites. In the case of F-actin ring formation, two distinct peaks are detected, correlating to the F-actin ring at the TJ (red arrowhead) and the posterior pole of the parasite (orange arrow). In the case of no F-actin ring, no accumulation at the TJ can be observed; only the peak correlating to posterior F-actin can be detected. For *adf* cKD, only one peak can be seen corresponding to the posterior pole of the parasite. **C.** Quantification of F-actin accumulation at the TJ for RH Cb-EmeraldFP parasites as detected in time lapse analysis. The values are expressed as percentage. 28 total parasite invasion events were captured across three different biological replicates. **D.** Representative immunofluorescence assays on indicated parasite strains during host cell penetration. While in the majority of control parasites F-actin can be detected at the junction (see quantification in (E)), no junctional ring can be detected for *adf c*KD, *act1 c*KO or *myoA* KO. Parasites were labelled with indicated antibodies. SAG1 staining was performed prior to permeabilisation to stain only the extracellular portion of the parasite. Antibodies against Ron2 were used to stain the TJ. Red arrowhead: position of the TJ. Orange arrow: posterior pole of the parasite. **E.** Tight Junction assay on fixed samples (as shown in (D)) depicting invasion rates of indicated parasites. Note that the number of total invasion events for *adf* cKD*, act1*cKO and *myoA*KO correlates to the number of invasion events, seen for RH without F-actin ring formation. **F.** Quantification of posterior F-actin accumulation during invasion of indicated parasite lines. Three biological replicates were done with 300 parasites counted for each biological replicate. One-way ANOVA analysis was performed for each graph. p value <0.0001. White arrows point to direction of invasion. Scale bars represent 5 µm.

Although we were unable to accurately quantify F-actin accumulation at the junction, we consistently observed that it increased during the invasion event (Fig.3A, MovieS6). At the onset of invasion, only a relatively faint F-actin signal could be detected at the apical tip of the parasite, its intensity increasing during invasion, culminating when the parasite invaded to ∼2/3. Collectively, while these observations fit well with the contribution of F-actin to the traction force applied at the TJ, the amount of invasion for which we could not detect F-actin at the TJ (25% show no F-actin ring) equalled the fraction of residual invasiveness detected for mutants for the acto-myosin system, such as shown in independent studies for mutants for *act1*, *mlc1, myoA, myoH, GAP45, adf, etc*. (Frenal, Polonais et al., 2010, Frenal & Soldati-Favre, 2015, Plattner, Yarovinsky et al., 2008, Tosetti et al., 2019) (Egarter et al., 2014, Meissner, Schluter et al., 2002, Whitelaw et al., 2017) (Mehta & Sibley, 2011)

While time lapse analysis of invasion events allowed documenting the dynamics of F-actin at the junction and posterior pole of the parasite, this analysis remained limited by insufficient resolution. Therefore, we decided to obtain higher resolution images using fixed assays to confirm the observed variability in F-actin accumulation at the junction. We quantified F-actin at the junction and the posterior pole using a modified red-green assay based on differential immunolabelling (Huynh & Carruthers, 2006). Invading parasites expressing Cb-EmeraldFP were labelled with SAG1 (non-invaded part of the parasite) and Ron2 (TJ-marker). We confirmed that in all cases parasites entered the host cell via a TJ, as indicated by positive Ron2 staining (Fig.3D).

In good correlation with live imaging analysis, we detected a F-actin ring at the TJ in ∼80% of RH-parasites, leaving ∼20% of the events where the presence of an F-actin ring remained questionable in fixed samples (Fig.3D,E). In contrast, in all invading parasites a posterior accumulation of F-actin was apparent (orange arrow, Fig.3D,F). In the case of *adf* cKD, which showed a reduced invasion rate of ∼20% compared to WT parasites, no F-actin ring at the junction was obvious in any invasion event (red arrowhead, *adf* cKD panel, Fig. 3D,E), while the posterior accumulation was still maintained (orange arrow, Fig.3D,F). As expected, no F-actin at the junction or at the posterior pole of the parasite was detectable in *act1* cKO parasites and only a cytosolic signal for Cb-EmeraldFP was apparent (Fig.3A,D,E,F). Similar to *adf*cKD an overall invasion rate of ∼20% was detected for *act1*cKO parasites, as described previously (Whitelaw et al., 2017), despite the posterior, aberrant morphology described previously (Egarter et al., 2014, Whitelaw et al., 2017) (green arrow, Fig.3D). Interestingly, in the case of *myoA*KO parasites, we found (similar to *adf*KD *and act1*cKO) an invasion rate of ∼20%, confirming previous results (Andenmatten et al., 2013, Whitelaw et al., 2017) (Meissner et al., 2002). However, disruption of MyoA resulted in loss of F-actin accumulation at both the junction site and the posterior pole, demonstrating that MyoA is the main motor required for retrograde F-actin flow during host cell invasion. Interestingly detectable F-actin appears to be equally distributed within the cytosol of the parasite and at the periphery of the parasite (Fig.3D,E,F), confirming the data obtained for extracellular parasites (Fig.1).

Together these data confirm that an intact acto-myosin system is required for efficient host cell invasion and that accumulation of F-actin at the posterior pole can occur independently of its accumulation at the junction, suggesting two independent processes, both dependent on MyoA-motor activity and F-actin dynamics. In analogy to other eukaryotes migrating through a constricted space, where the nucleus represents a major obstacle that needs to be deformed, squeezed and pushed through the constriction (Thiam et al., 2016), we speculated that the nucleus and potentially other bulky organelles of the parasite represent a major limitation for invasion that need to be forced through the junction.

### Posterior localised F-actin seems to push the nucleus through the junction

We next simultaneously visualised F-actin dynamics and the position and shape of the nucleus using time lapse imaging (Fig.4). We confirmed that the accumulation of F-actin at the junction became most prominent, once the nucleus reached the junction (orange arrow, Fig.4A, Movie S7). Interestingly, in the vicinity of F-actin located at the parasite posterior pole we observed a cup-like meshwork around the nucleus closed to the junctional F-actin ring (blue arrow, Fig.4A, Movie S7). Once the nucleus reached the junction it became constricted and deformed when the F-actin ring is visible, (see red arrowhead 0:15-0.21, Fig.4A.B). When no F-actin ring was detectable (Fig.4B), the shape of the nucleus remained relatively constant (red arrows, Fig.4D,E). However, independently of F-actin ring formation, the posterior pole of the parasite contracted (blue arrows in Fig.4A.D; Movie S7) prior or coinciding with nuclear entry, which was also confirmed by measuring the diameter of the posterior pole of the parasite during the invasion process (Fig.4C,F).

**Figure 4.**
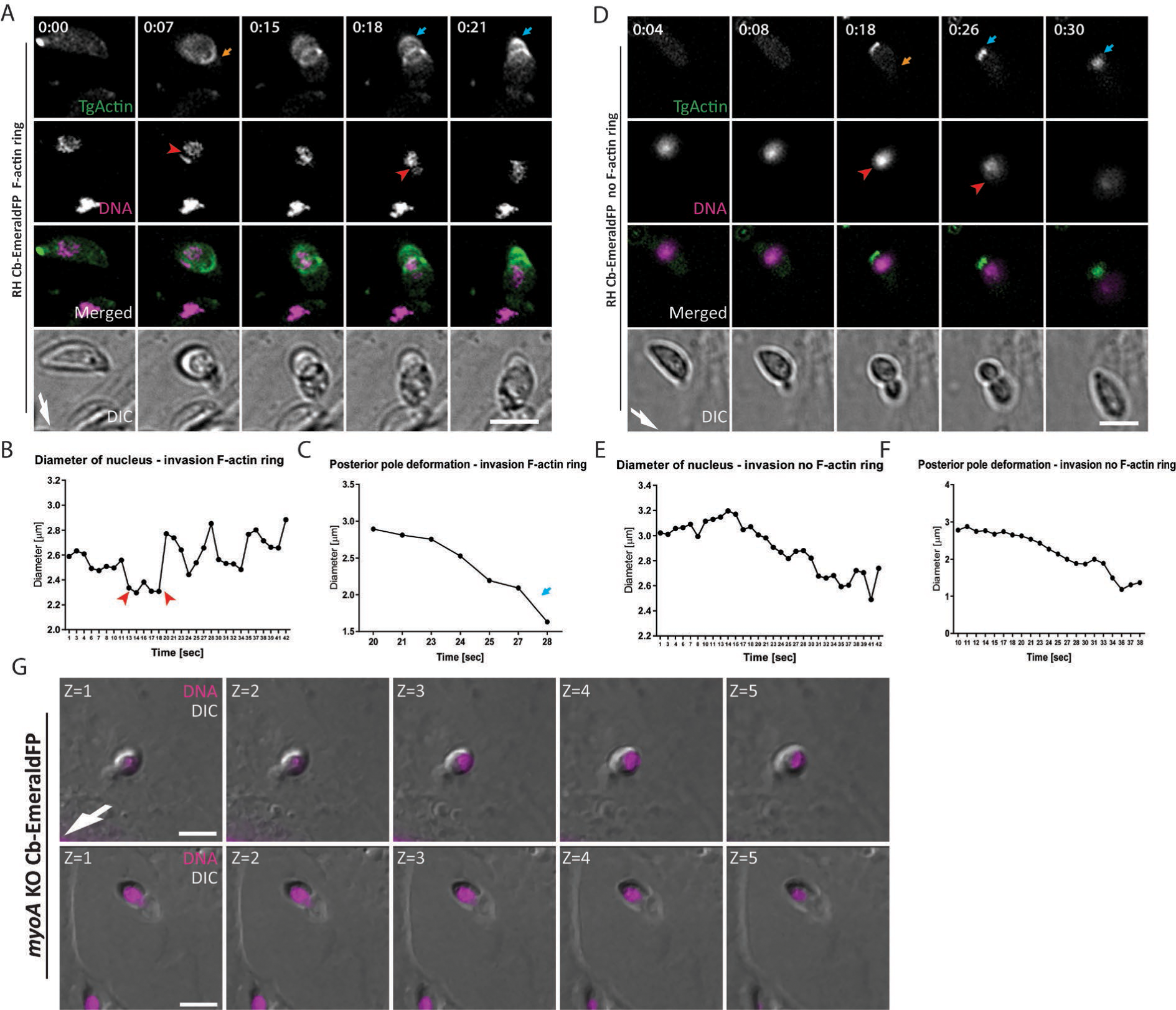
Time lapse analysis of F-actin and nuclear dynamics during host cell invasion **A.** Time-lapse analysis of invading RH Cb-EmeraldFP parasites. During penetration, an F-actin ring is formed at the TJ. The nucleus (purple) is squeezed through the TJ (red arrowhead) and posterior F-actin appears to be directly connected to the nucleus. **B.** Analysis of nucleus deformation during live imaging shown in A. The nucleus constricts while its passing through the TJ (red arrowheads). **C.** The posterior end of the parasite deforms during the invasion process. F-actin is accumulated during invasion, with the posterior end contracting. **D.** Time-lapse stills depicting RH Cb-EmeraldFP parasites invading in absence of detectable F-actin at the junction. The nucleus (purple) is squeezed through the TJ (red arrowhead) during invasion. Actin accumulation at the posterior pole of the parasite is strongly detected in all cases (blue arrow). **E.** Analysis of nucleus deformation during live imaging shown in (D). The nucleus appears to not show constriction once its passing through the TJ when the F-actin ring is not present. **F.** The posterior end of the parasite also deforms during the invasion process. **G.** Z-stack gallery depicting mid-invading MyoAKO parasites. Note that the nucleus is located located at the back during invasion events. Scale bar represents 5 µm. Numbers indicate minutes:seconds. Scale bar represents 5 µm. White arrow points to direction of invasion.

We then imaged the nucleus of *myoA*KO parasites over their residual host cell invasion (Fig.4G). Since the MyoA-motor-complex is known to generate the required traction force at the TJ (Bichet et al., 2016), we interpret these data as if optimal entry by WT parasites results from a coordinated mechanism, requiring the glideosome, acting below the surface of the parasite(Tardieux & Baum, 2016), together with cytosolic localised F-actin accumulating at the posterior pole and surrounding the nucleus. In good support, when no F-actin was detectable at the junction, indicating independence of traction force generated at the junction, posterior accumulation of F-actin adjacent to the nucleus was still seen during invasion (Fig.4D-F, MovieS7).

### F-actin forms a continuous network between the junction and posterior accumulation that surrounds the nucleus

We applied super-resolution structure illumination microscopy (SR-SIM) to analyse the distribution of F-actin in WT parasites that were at the mid of invasion and found consistently that the nucleus of the parasite is surrounded by F-actin that formed a continuous meshwork connected to the posterior pole of the parasite (Fig.5A-D). Importantly, this meshwork could be observed independently of F-actin accumulation at the junction but broke down upon disruption of *adf* and was significantly reduced in the case of *myoA*KO parasites (Fig.5A,D Movie S8). The perinuclear F-actin meshwork formed a continuum with the posterior accumulation (Fig.5B). Finally, 3D-models demonstrated that F-actin is perinuclear and can be observed in ∼60% of all cases in WT parasites, while it is significantly reduced in *myoA*KO (∼20%) and absent in *adf*KD parasites (Fig.5D).

**Figure 5.**
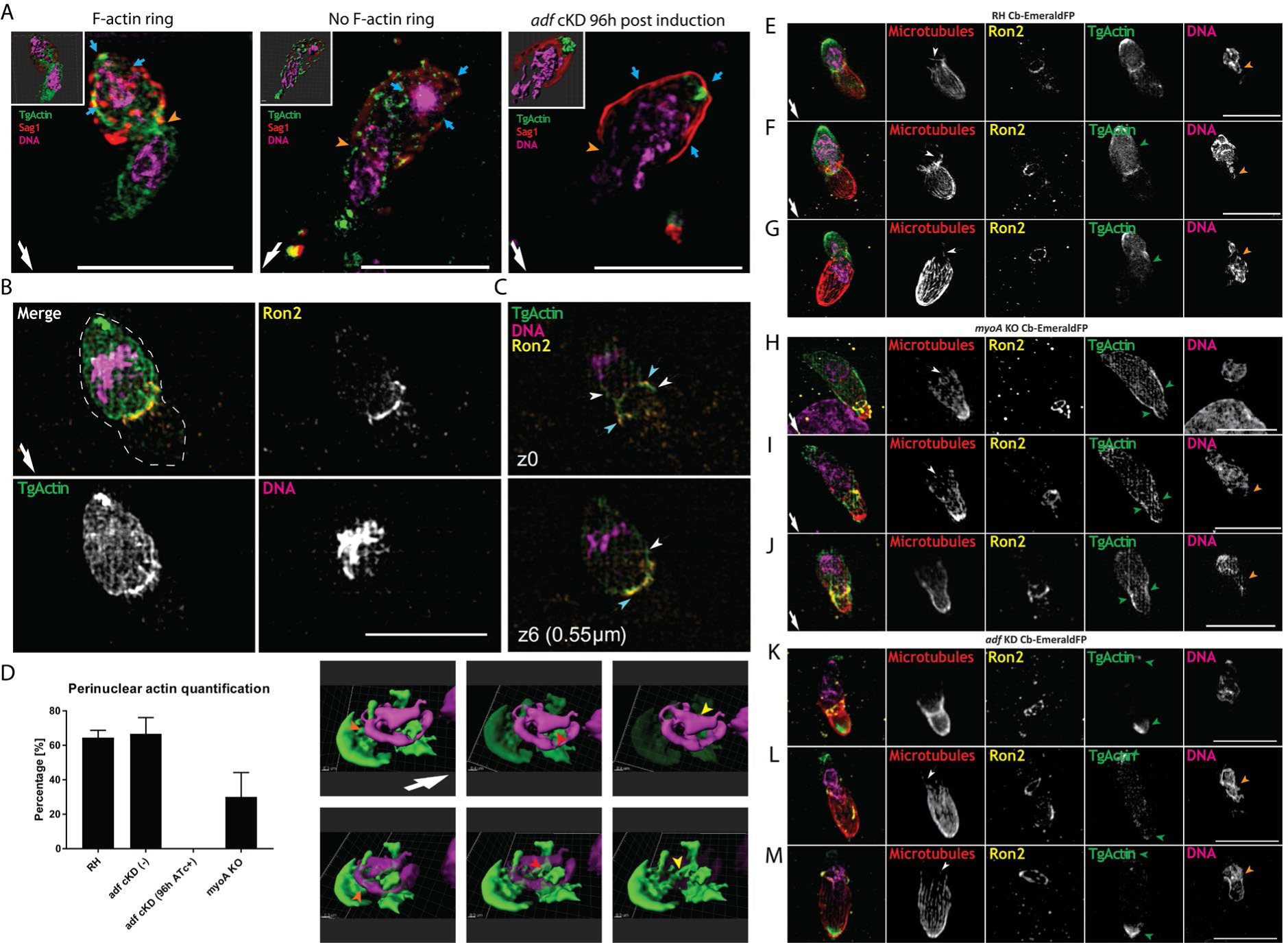
Super-resolution microscopy demonstrates formation of a F-actin cage around the nucleus during invasion. **A.** SR-SIM images depicting invading parasites with and without the presence of the F-actin ring at the tight junction and invading *adf* cKD Cb-EmeraldFP parasites. SAG1 staining (in red) was performed prior to permeabilisation to specifically label extracellular part of the parasite. Note the accumulation of actin (blue arrow) around the nucleus and the posterior pole of the parasite, irrespective of F-actin ring formation at the TJ (orange arrowhead). Imaris 3D rendering are shown for each presented case (inlet). Scale bar represents 5 µm. White arrow indicates direction of invasion. **B.** SR image of an invading parasite before nuclear entry. F-actin forms a continuous meshwork around the nucleus that appears connected to the posterior pole. **C.** F-actin localisation in relation to the TJ during invasion. F-actin is in close vicinity to the Ron2 ring during invasion. **D.** Quantification of perinuclear actin in indicated parasites. Parasites were counted by assessing Cb EmeraldFP signal as an actin marker around or in the vicinity of the nucleus. Still images to the right represent what was considered as F-actin – nucleus association. The orange arrowhead points to F-actin surrounding the nucleus in the posterior end of the parasite, the red and yellow arrowhead shows F-actin in the vicinity of the nucleus, hinting at a close interaction between the two. **E-G.** SR-SIM images showing three stages of invading wt parasites. SiR-tubulin staining for microtubules (in red) was performed prior to fixation to specifically label microtubules in the parasite. During invasion, the microtubules (MT) are deformed at the TJ area. Accumulation of actin (green arrow) at the posterior forms a F-actin meshwork. The nucleus is deformed when it enters the TJ area (orange arrowhead). **H-J.** In the case of the *myoA* KO strain, the MTs also deform when passing through the TJ. However, F-actin is not concentrated at the posterior pole or the TJ, but evenly distributed within the cytosol of the parasite, with some peripheral location. **K-M.** In case of *adf*KD, MTs are deformed at the TJ area. F-actin is accumulated at both ends of the parasite (green arrowheads), with no accumulation at the TJ. Scale bar represents 5 µm. White arrow points to direction of invasion.

In most migrating cells the nucleus is positioned in the back, which requires the coordinated action of posterior F-actin as well as anterior microtubules, resulting in positioning of the nucleus by a pull-and-push mechanism driven by the interplay of an actomyosin complex, actin-nucleus interface proteins and other cytoskeletal structures (McGregor et al., 2016). In the case of *T.gondii* the subpellicular microtubules are polymerised from microtubule organising centres at the apical tip, aligned on the cytosolic side of the IMC, where they directly interact with gliding associated membrane proteins (GAPMs; (Harding et al., 2019)). This subpellicular microtubules form a basket that covers ∼2/3 of the parasite length. Furthermore, it was previously suggested that parasite F-actin can interact with the subpellicular microtubules (Patron et al., 2005) (Yasuda et al., 1988), leading to the question if the basally localised meshwork consisting of F-actin might interact with the apically localised subpellicular microtubules during invasion. Therefore, we assessed the location of microtubules (MTs) during invasion by the use of SiR-tubulin. The MTs covers 2/3 of the parasite body creating a rigid structure starting from the apical tip (Fig.5E-M; MovieS9). During invasion, this MT structure appears to be significantly deformed, with several examples detected, where MTs seem to colocalise with junctional F-actin (white arrows in Fig.5). Furthermore, MTs appear frequently bent and distorted at the junction. Interestingly, this polarisation with F-actin at the posterior end and microtubules at the leading edge is maintained throughout the invasion process, from early to late stage (Fig.5, Fig.S6). In some cases, the MTs extended far beyond the 2/3 parasite body, aligning the nucleus (white arrow, Fig.5). The microtubules closely align to the part of the nucleus that enters through the TJ (orange arrowheads), while F-actin closely aligns to the posterior part of the nucleus (green arrowheads), resulting in a highly polarised cytoskeletal organisation around the nucleus (Movie S9). It is plausible that akin to other eukaryotes, a coordinated action of microtubule and F-actin based mechanisms supports nuclear entry and deformation through the TJ. This basic organisation is also maintained in those cases, where no F-actin accumulation is detectable at the junction (Fig.5G).

Upon interference with actin dynamics (*adf*cKO) or the glideosome (*myoA*KO) this organisation is disrupted. In case of ADF depletion, posterior accumulation of F-actin is still apparent (green arrowheads, Fig.5K-M), but no F-actin meshwork around the nucleus, connecting to subpellicular microtubules is detectable (orange arrowheads, Fig.5K-M). In contrast, disruption of MyoA results in a dominant arrangement of F-actin throughout the cytosol with no accumulation at the posterior pole of the parasite or formation of perinuclear F-actin (Fig.5H-J), suggesting that there is a coordination of glideosome function in between the PM and IMC and the formation of the cytosolic localised meshwork around the nucleus. It is also noteworthy, that F-actin accumulates in the region of the TJ (green arrowheads) when compared to the rest of the cell.

In summary, the combined data from fixed invasion assays and SR-SIM imaging allow documenting that the parasite’ nucleus is surrounded by a F-actin meshwork and that its formation and coordination might depend on subpellicular microtubules. Here filaments appear to adapt a cytoskeletal conformation designed to protect, deform and push the nucleus inside the host cell and we propose that during invasion the glideosome provides the traction force for general movement of the parasite through the TJ, while perinuclear actin is important in order to facilitate nuclear entry.

### Perinuclear F-actin is detected during invasion of Plasmodium merozoites

While *T. gondii* is a relevant model to address general questions in apicomplexan biology, we previously observed differences in the role of F-actin during the asexual life cycles of *T. gondii* and *P. falciparum*, the causative agent of human malaria. For instance, in case of *P. falciparum*, invasion is dependent on parasite F-actin (Das, Lemgruber et al., 2017), while in the case of *T. gondii* residual invasion can be observed. To analyse the role of F-actin during merozoite invasion, we expressed CB-Halo and Cb-EmeraldFP in *P. falciparum* and validated their efficiency to analyse F-actin localisation and dynamics in this parasite (Stortz et al., BioRXiv2019). Invasion assays and localisation analysis of F-actin during merozoite invasion showed a similar distribution of F-actin at the junction, around the nucleus and at the posterior pole of the parasite (Fig.6A, C). We also analysed F-actin localisation during the invasion of *P. knowlesi* merozoites using a previously described antibody that preferentially recognises *P.falciparum* F-actin and obtained identical results(Angrisano et al., 2012). Fixed immunofluorescence imaging of merozoite’s in the process of invasion were analysed for the distribution of F-actin. As with *T. gondii*, F-actin can be detected at the junction, at the onset of invasion (Fig.6) pointing to a similar strategy between apicomplexan parasites. Importantly, super-resolution imaging of F-actin in merozoites in different stages of invasion, clearly highlight accumulation of F-actin at the posterior pole and in direct contact with the nucleus (Fig.6C).

**Figure 6.**
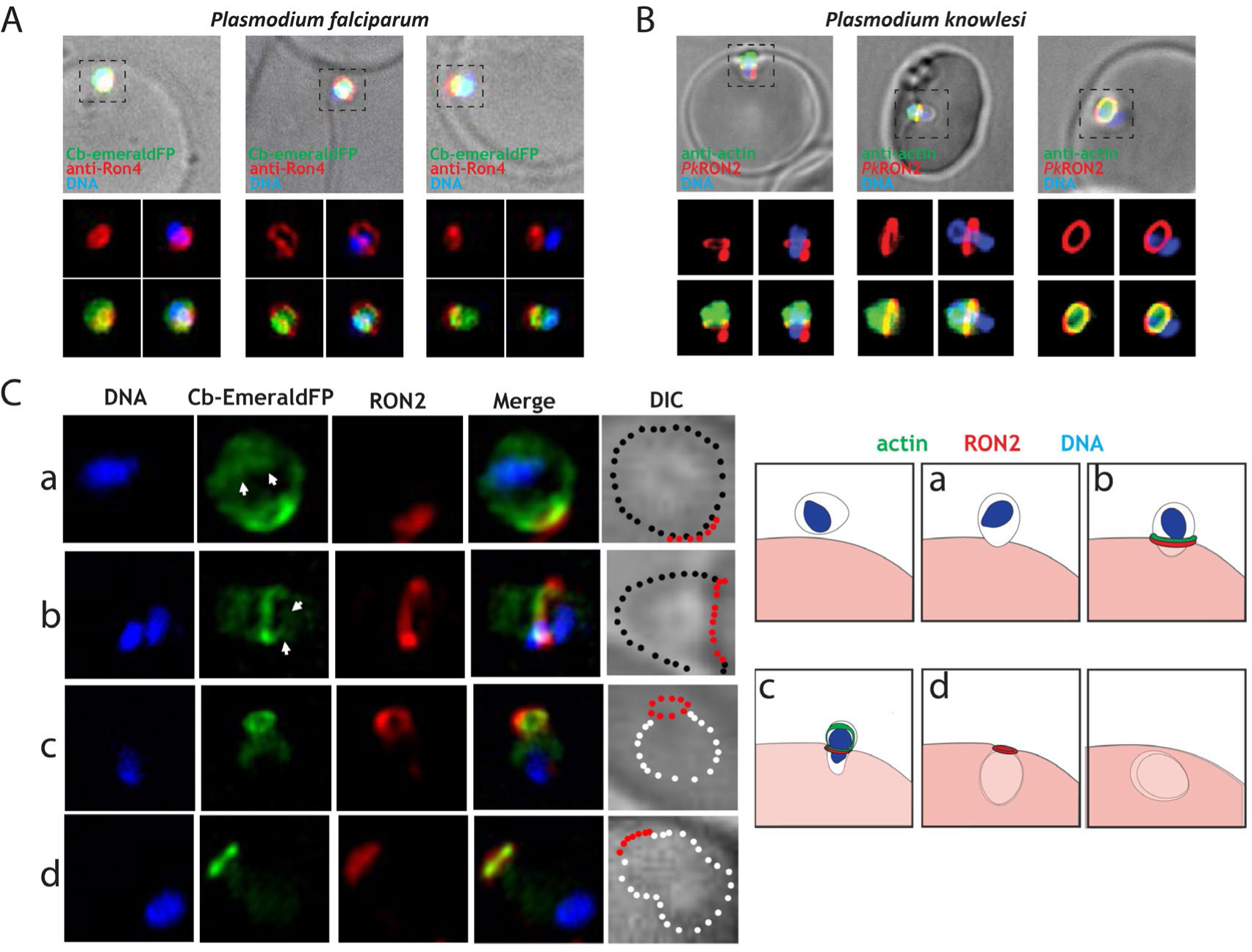
*Plasmodium* F-actin localisation during invasion. **A.** IFA showing various stages of invasion of erythrocytes by *P. falciparum* merozoites expressing Cb-EmeraldFP. DAPI labels the nucleus (DNA), Cb-EmeraldFP labels actin filaments (CB-EME, green) and the invasion junction is marked with an anti-RON4 antibody (red). Left panel: a merozoite beginning invasion; note the formation of an F-actin ring (green) behind the RON4 junction (red). Middle panel: A merozoite in the middle of the invasion event; note the constriction of the nucleus (blue) as it passes through the RON4 junction and F-actin colocalising with it (green). Right panel: A merozoite completing the process of invasion; note the F-actin ring beyond the RON4-junction and actin filaments surrounding the nucleus. **B.** IFA showing various stages of invasion of erythrocytes by *P. knowlesi* merozoites. DAPI labels the nucleus (DNA), anti-actin Antibody labels actin filaments (anti-actin, green) and the invasion junction is marked with a C-terminally tagged *Pk*RON2 mCherryHA (red). Left panel: a merozoite beginning invasion; note the formation of an F-actin ring (green) behind the *Pk*RON2 junction (red). Middle panel: A merozoite in the middle of the invasion event; note the constriction of the nucleus (blue) as it passes through the *Pk*RON2 junction and F-actin colocalising with it (green). Right panel: A merozoite completing the process of invasion; note the F-actin ring beyond the *Pk*RON2-junction and actin filaments colocalising with the nucleus. **C.** Super-resolution images showing various stages of invasion of erythrocytes by *P. falciparum* merozoites expressing Cb-EmeraldFP. DAPI labels the nucleus (blue), Cb-EmeraldFP labels actin filaments (CB-EME, green) and the invasion junction is marked with an anti-RON4 antibody (red). Brightfield images have been marked with red dots depicting the junction, black dots depicting merozoite boundary still outside the erythrocyte and white dots depicting a merozoite that has penetrated the host cell. Top panel shows an attached merozoite beginning the process of invasion, the second panel during the process of invasion and the bottom two panels show merozoites completing the process of invasion. Right panel: Schematic depicting merozoite invasion into red blood cells. Letters depict the corresponding event in the model (left) and microscopy pictures (right). White arrows depict actin filaments that colocalise with the nucleus.

## Discussion

Host cell invasion by apicomplexan parasites occurs through a tight junctional ring (TJ), established by the parasite upon secretion of its rhoptry content (Besteiro et al., 2011). Previous studies demonstrated that the TJ is anchored to the host cell cortex via its interaction with host cell factors (Gonzalez, Combe et al., 2009, Guerin et al., 2017) thereby acting as an anchor for traction force exerted by the parasites acto-myosin-system (Bichet et al., 2016). In addition, compressive forces, caused by the host cell at the TJ requires the deformation of the parasite during invasion, which appears to be counterbalanced to ensure integrity of the parasite (Bichet et al., 2016). In fact, the invasion of host cells through the TJ appears in many aspects very similar to the migration of eukaryotic cells through a constricted environment, with impressive morphological changes observed at the point of constriction (Petrie & Yamada, 2016). Here, the nucleus represents a physical barrier for migration that requires to be modulated, deformed and protected in order to be squeezed through the constriction (Thiam et al., 2016). During migration through a constricted space, rapid actin nucleation around the nucleus facilitates nuclear deformation and protection. Furthermore, proteins that interface actin and the nucleus such as the family of LINC (Linker of Nucleoskeleton and Cytoskeleton) proteins allow the formation of intricate perinuclear F-actin structures connected to microtubules and other cytoskeletal elements that support nucleus translocation, mechanotransduction and protection (Guilluy & Burridge, 2015, Guilluy, Osborne et al., 2014).

In apicomplexan parasites, a similar function of F-actin was initially ruled out based on the assumption that F-actin acts exclusively within the narrow space between the PM and IMC, where the glideosome is localised (Frenal et al., 2017). However, the finding that the majority of F-actin dynamics appears to occur within the cytosol of the parasite in extracellular parasites, and the fact that host cell invasion is significantly slowed down once the parasites nucleus reaches the TJ and is usually blocked at this stage in the case of mutants for the acto-myosin system, led us to the hypothesis that parasite F-actin serves a second role during invasion, namely the facilitation of nuclear entry, akin to other eukaryotes, where a squish and squeeze model has been proposed (McGregor et al., 2016).

To test this hypothesis, we performed a careful analysis of WT parasites and mutants for core components of the acto-myosin system and correlated their phenotypes with the observed changes in F-actin dynamics in extracellular and invading parasites, using quantitative imaging approaches allowing us to describe the dynamics of F-actin during host cell invasion. We used a combination of real-time and super-resolution microscopy to demonstrate that the nucleus is indeed a major obstacle that needs to be overcome for successful invasion. In good agreement, mutants of the acto-myosin system demonstrate an extended pause during invasion, as soon as the nucleus reaches the junction. In all cases, parasites accumulate F-actin at the posterior pole during invasion, which forms a continuous meshwork with perinuclear F-actin. In contrast, F-actin accumulation at the junction is not always observed during the invasion process, suggesting that the glideosome activity is fine-tuned, depending on the force required for host cell penetration. Importantly, F-actin accumulation at the posterior pole can occur independently from F-actin accumulation at the junction, suggesting the action of two independent pools of F-actin that contribute to invasion and nuclear entry. Finally, we demonstrate that parasite F-actin and subpellicular microtubules form a highly polarised cell during invasion, with the subpellicular microtubules at the leading edge and F-actin at the posterior pole, suggesting tight and controlled coordination of these two cytoskeletal elements to facilitate host cell invasion.

While the interplay between the F-actin and microtubule cytoskeleton of the parasite needs to be further analysed, it is interesting to note that deletion of MyoA, the core motor of the glideosome (Meissner et al., 2002) results in complete loss of F-actin polarity and an even distribution of F-actin in the cytosol of the parasite, with some peripheral accumulation during motility and invasion. This indicates a tight interlinkage of F-actin dynamics potentially occurring in between the PM and IMC with the F-actin dynamics observed within the cytosol of the parasite. This is also reflected, by the fact that F-actin dynamics originating from the apical tip of the parasite, potentially by Formin-1 (Tosetti et al., 2019) and close to the Golgi by Formin-2 (Tosetti et al., 2019)(Stortz et al., BioRxiv2018) forms a continuous network that appears to be interlinked as presented in the 3D-SIM of this work. Interestingly, stimulation of Calcium-signalling with either A23187 or BIPPO resulted in preferential F-actin flow towards the basal end of the parasite. Surprisingly, F-actin dynamics in unstimulated *myoA*KO parasites is unchanged, but upon stimulation F-actin relocation appears to be blocked and it remains “stuck” in the first half of the parasite, demonstrating that the glideosome is required for efficient relocation of the whole F-actin pool and that both processes are tightly interlinked. Analysis of host cell invasion demonstrated that F-actin can be found at the TJ in most cases and -as expected-disruption of MyoA, ADF or Act1 results in highly reduced invasion rates, where no F-actin can be detected at the TJ. It remains to be seen, if the F-actin ring is formed in order to stabilise the junction to counteract pressure exerted by the host cell (Bichet et al., 2016) or if the activity of the glideosome can be modulated, depending on the necessity to provide the traction force to facilitate entry of the parasite. Additionally, another question remains on how the parasite regulates the formation and translocation of this F-actin ring, as unlike a report where FRM1 translocates along the TJ in *P.falciparum* during invasion (Baum et al., 2008), *T.gondii* did not present a similar mechanism (Jacot et al., 2016). Collectively, while these observations fit well with the contribution of F-actin to traction force applied at the TJ, the amount of invasion for which we could not detect F-actin at the TJ (25% show no F-actin ring) equalled the fraction of residual invasiveness detected for mutants for the acto-myosin system, such as shown in independent studies for mutants for *act1*, *mlc1, myoA, myoH, GAP45, adf, etc*. (Frenal et al., 2010, Frenal & Soldati-Favre, 2015, Plattner et al., 2008, Tosetti et al., 2019) (Egarter et al., 2014, Meissner et al., 2002, Whitelaw et al., 2017) (Mehta & Sibley, 2011).

A similar situation is seen for the formation of perinuclear actin during host cell invasion, with most parasites demonstrating the formation of an F-actin meshwork around the nucleus. Again, disruption of the acto-myosin system causes disruption of this meshwork, demonstrating a tight interplay between glideosome function and formation of perinuclear actin. In contrast, in all cases F-actin is seen to accumulate at the basal pole of the parasite during invasion, which is connected to the perinuclear meshwork (if formed during invasion). Finally, live imaging analysis indicates that the basal end of the parasite contracts during host cell entry, potentially forcing the nucleus through the junction.

In most migrating cells, the nucleus is positioned in the back, which requires the coordinated action of posterior F-actin as well as anterior microtubules, resulting in positioning of the nucleus by a pull-and-push mechanism driven by the interplay of an actomyosin complex, actin-nucleus interface proteins and other cytoskeletal structures such as intermediate filaments and microtubules (McGregor et al., 2016). In the case of *T.gondii*, the subpellicular microtubules are polymerised from microtubule organising centres at the apical tip, aligned on the cytosolic side of the IMC, where they directly interact with components of the glideosome, such as the gliding associated membrane proteins (GAPMs; (Harding et al., 2019)). Indeed, it was previously suggested that parasite F-actin can interact with the subpellicular microtubules (Patron et al., 2005) (Yasuda et al., 1988), leading to the question if the posterior localised meshwork consisting of F-actin might interact with the apically localised subpellicular microtubules during invasion to form a highly polarised cellular network, as seen in Figure 5 (S5). Further studies are required to analyse the linkage of these two cytoskeletal elements as subpellicular microtubules could function as stabilisers of cell shape during invasion, allowing the coordination of the forces required for host cell invasion.

Together these results permit us, for the first time, to reconcile discrepancies in the phenotypic invasion characteristics of apicomplexan parasites with F-actin imaging data and to propose a new model for actin’s contribution during host cell invasion, in particular nuclear entry through the TJ (Fig.7). While the IMC-localised acto-myosin motor complex provides traction force at the TJ to initiate and facilitate cell invasion, the entry of the parasite nucleus into the host cell appears to be a major limiting step that requires direct action of a cytosolic acto-myosin system, possibly also involving the subpellicular microtubules in order to be deformed, protected and pushed into the host cell (Fig.7).

**Figure 7.**
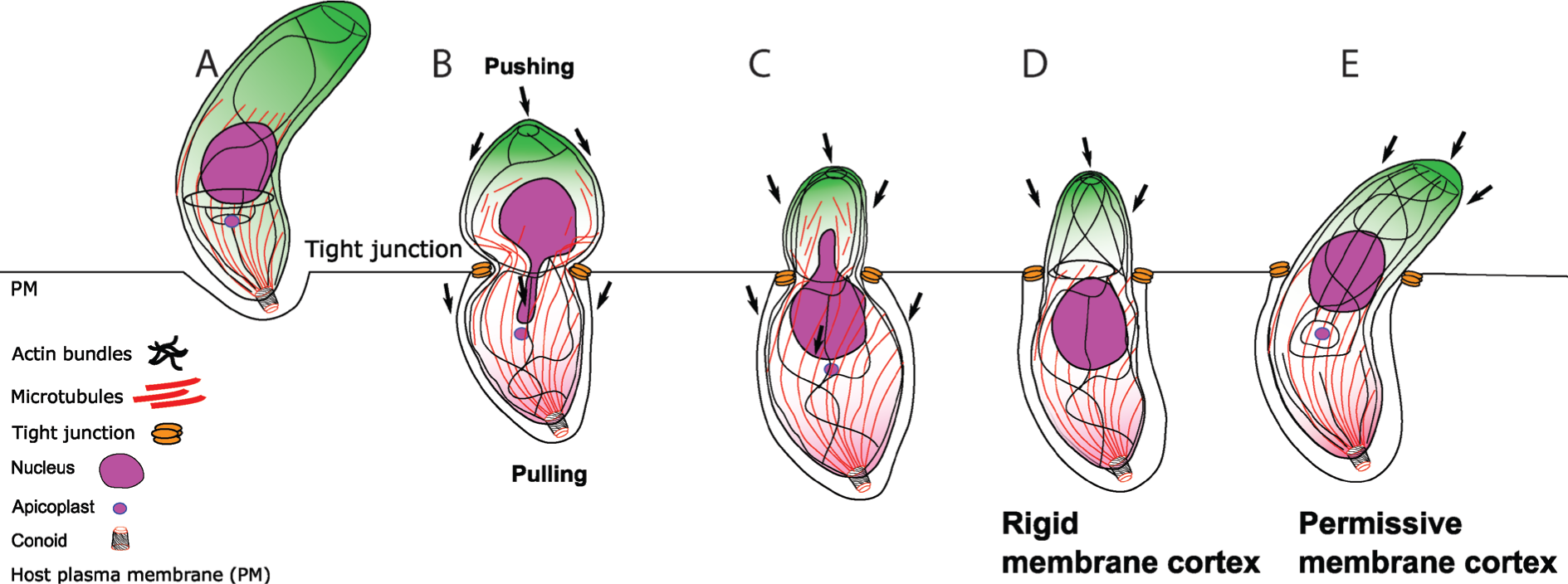
Model of the proposed nuclear squeeze mechanism during apicomplexan invasion. **A-D.** Once the parasite attaches to the surface of a host cell, F-actin strongly accumulates at the posterior pole and at the apical end. During penetration, the junctional complex is formed that contributes to the attachment and stabilisation of the parasite to the host cell in the TJ. F-actin at the TJ provides traction force and stability for nuclear entry, while posterior F-actin provides contraction force to allow nuclear entry. At the same time microtubules might facilitate nuclear entry by pulling. We propose that the nucleus is squeezed through the TJ by a pushing-pulling mechanism controlled by actin and potentially microtubules. **E.** In some cases (for example due to more permissive host cells or upon modulation of F-actin dynamics), a F-actin ring at the junction is not required/formed and the nucleus can enter the host cell by the action of posterior accumulated F-actin.

## Material and Methods

### Plasmid construction

The same procedure was carried out as described previously^12^. The Cb-EmeraldFP plasmid consists of a sequence encoding actin chromobody (Cb) from Chromotek followed downstream by an in frame sequence encoding EmeraldFP. The vector backbone contains a *DHFR* promoter for protein expression without drug resistance genes. To create Cb-EmeraldFP, the EmeraldFP coding sequence was amplified using primers (EmeraldFP-F: atgcaccggtatgggactcgtgagcaaggg and EmeraldFP-R: atgccttaagttacttgtacagctcgtcca). The EmeraldFP PCR product plasmid were subcloned using traditional restriction digestion/ligation protocols.

### Culturing of parasites and host cells

Human foreskin fibroblasts (HFFs) (RRID: CVCL_3285, ATCC) were grown on tissue culture-treated plastics and maintained in Dulbecco’s modified Eagle’s medium (DMEM) supplemented with 10% foetal bovine serum, 2 mM L-glutamine and 25 mg/mL gentamycin. Parasites were cultured on HFFs and maintained at 37° C and 5% CO2. Cultured cells and parasites were regularly screened against mycoplasma contamination using the LookOut Mycoplasma detection kit (Sigma) and cured with Mycoplasma Removal Agent (Bio-Rad) if necessary.

### *T. gondii* transfection and selection

To generate stable Cb-EmeraldFP expressing parasites, 1 × 10^7^ of freshly released RH Δ*hxgprt*parasites and *adf* cKO were transfected with 20 µg DNA by AMAXA electroporation. Transfected parasites were then sorted using flow cytometry with a S3 Cell Sorter (Bio-Rad, Hercules, CA, USA).

### *Merozoite* isolation

PkRON2*mCherry-HA A1-H.1 merozoites were isolated as described^21^. In brief, ring-stage parasites were tightly synchronised to a 3.5 hrs and allowed to develop to schizont stage. Schizonts were separated from uninfected erythrocytes by MACS magnet separation (CS column, Miltenyi Biotec). Parasites were returned to complete media at 37 oC and incubated with 10 µM E64 (Sigma-Aldrich) for <4 hrs. Schizonts were pelleted (850 × g, 5 mins) and resuspended in incomplete media and merozoites purified by 3 µm filtration (Whatman) immediately followed by 2 µm filtration (Whatman).

### Inducing the conditional LoxP*act1* cKO

The inducible *act1* cKO was obtained by the addition of 50 nM rapamycin to the parental LoxPAct1 strain for 4 hr at 37°C, 5% CO_2_ and cultured as described previously^6^.

### Inducing the conditional Cas9*act1* cKO

The act1 cKO parasite strain was obtained by addition of 50 nM of rapamycin to the parental lines. The strain was incubated for 1 h at 37 °C and 5% CO2 and cultured as described previously^6^. To decrease the population of un-induced parasites, the culture media was replaced by DMEM complete supplemented with 2.5% dextran sulphate after 24 h to inhibit re-invasion of WT parasites.

### Light microscopy

Widefield images were acquired in z-stacks of 2 μm increments and were collected using an Olympus UPLSAPO 100× oil (1.40NA) objective on a Delta Vision Core microscope (AppliedPrecision, GE) attached to a CoolSNAP HQ2 CCD camera. Deconvolution was performed using SoftWoRx Suite 2.0 (AppliedPrecision, GE). Video microscopy was conducted with the Delta Vision Core microscope as above. Normal growth conditions were maintained throughout the experiment (37° C; 5% CO_2_). Further image processing was performed using FIJI for ImageJ (NIH) and Icy Image Processing Software (Institut Pasteur) ^22^. Super-resolution microscopy (SR-SIM) was carried out using an ELYRA PS.1 microscope (Zeiss) as described previously^12^. 3D rendering and models views were generated using Imaris software (Bitplane, Oxford Instruments) using the acquired SR-SIM files.

### Flow analysis time-lapse microscopy

To discriminate parasite orientation, 500nm of SiR tubulin was added to parasite cultures 1 hour before the experiment. To prepare tachyzoites for the assay, 1 × 10^6^ mechanically freshly released parasites per live dish were washed 3 times with PBS and incubated a minimum of 10 minutes in serum-free DMEM media. Cytochalasin D was added 30 minutes before imaging in a concentration of 2 µM. The samples were then imaged by widefield microscopy using a Delta Vision Core microscope or Super-resolution microscopy (SR-SIM) using an ELYRA PS.1 microscope.

### Parasite treatment with A23187 and BIPPO

Calcium ionophore A23187 was added to parasite live dishes using with a final concentration of 2µM. BIPPO was similarly used with a final concentration of 5µM. The parasites were imaged by light microscopy immediately after adding A23187 or BIPPO.

### Tight Junction assay

For the assay 3 × 10^6^ mechanically freshly released parasites per well were incubated in 200 µl Endo Buffer (44.7 mM K2SO4, 10 mM MgSO4, 106 mM sucrose, 5 mM glucose, 20 mM Tris– H2SO4, 3.5 mg/mL BSA, pH 8.2) for 10 minutes at 37° C degrees. The parasites were then allowed to settle into confluent layer of HFFs for 3 minutes at room temperature. This step was followed by 20 minutes incubation at 37° C. The supernatant was carefully removed and replaced by pre-warmed culture media. The samples were further incubated for 5 minutes to allow invasion.

This was followed by fixing the samples using two different buffers: Cytoskeleton buffer (CB1) (MES pH 6.1 10mM, KCL 138 mM, MgCl 3mM, EGTA 2mM, 5% PFA) and Cytoskeleton buffer (CB2) (MES pH6.1 10 mM, KCL 163.53 mM, MgCl 3.555 mM, EGTA 2.37 mM, Sucrose 292 mM). These buffers were mixed in a 4:1 ratio respectively and subsequently used for fixation for 10 minutes. The samples were then treated with a PFA quenching solution (NH_4_CL 50 mM) for 10 minutes followed by PBS washing for three times. A minimum total of 40 parasites were counted for each sample in three biological replicates. The tight junction assay involving several mutants and wt in Fig. 2d employed a multiple comparison 2way ANOVA using Tukey’s multiple comparison test with a p-value of <0.0001

### Cell deformation analysis

Samples were captured using SR-SIM and widefield microscopy as previously described^12^. The deformation was then analysed with Zen blue software (Zeiss) by measuring the size of the parasite body and nucleus when going through a tight junction. A total of 100 parasites per biological replicate were measured for each condition for RH in Fig. 1b and a total of 30 parasites per biological replicate were measure for the mutant in Fig. S1a. Statistical analysis was carried out by ordinary one-way ANOVA, using Tukey’s multiple comparison test with a p-value of <0.0001.

### Color-coded kymogram generation for particle dynamics analysis

Fourier-filtered Color-coded kymograms were generated using the KymographClear plugin on ImageJ and published in Mangeol, Prevo, & Peterman, 2016. This plugin allows to trace a path of interest on a collapse t-stack to follow particle movement. The plugin is able to generate a three color-coded kymogram using Fourier transformation to filter different populations of particles by the orientation of the trajectory. The color-coding is represented by forward movement as red, backward movement as green and no movement as blue. This data is then exported to the stand-alone software KymographDirect to stablish time-average intensity profiles. A more detailed protocol can be found on the author’s website: https://sites.google.com/site/kymographanalysis/

### Skeletonisation Analysis

The skeletonisation process was done in ImageJ FIJI using the Skeletonisation processing plugin(Arganda-Carreras et al., 2010). The sample was binarized by thresholding. A skeletonisation algorithm is them employed which converts the thresholded signal into pixels, reducing the total width but not the length. The time-lapse profile was collapsed (Z-Project) to define the F-actin structures and their dynamics over the course of the time-lapse. The plugin can be accessed by: https://imagej.net/AnalyzeSkeleton

### Live cell invasion

Parasites were mechanically released using 23 G needle and filtered prior to inoculation on a confluent layer of HFFs, grown on glass bottom dishes (MaTek). The dish was then transferred to the DV Core microscope (AppliedPrecision, GE) and maintained under standard culturing conditions. Images were captured at 1.4 frames per second in DIC using an Olympus apochromat 60x oil objective. Images were analysed using the Icy Image Processing software (Institut Pasteur)^5^. Penetration speed profiles were obtained for 18 independent invasion events for RH Cb-EmeraldFP and Cb-EmeraldFP parasites. The statistical analysis of RH wt Invasion events in Fig. 2c was done by unpaired two-tailed t-test in 28 total parasite invasion events that were captured across three different biological replicates.

### Penetration Speed profile analysis

Movies obtained were analysed using Icy Image Processing software (Institut Pasteur)^22^ with the Manual Tracking plugin. The methodology used consist on tracking the apical end of the parasite through the duration of penetration. The resulting data was then exported to GraphPad PRISM7 Software for posterior analysis.

### Correlative light-electron microscopy (CLEM)

Cells were grown in gridded glass bottom petri dishes (MaTek) and infected with Cb-EmeraldFP or RH parasites. Invading parasites were imaged with SR-SIM in an ELYRA PS.1 microscope (Zeiss, Germany) with a Plan-Apochromat 63x oil (1.4 NA) objective. After imaging, the material was fixed in 2.5% glutaraldehyde and 4% paraformaldehyde in 0.1 M Pipes buffer (TAAB, UK). To increase the contrast of cytoskeleton elements, 1% tannic acid was added to the fixation solution. After an overnight incubation, the material was processed for transmission electron microscopy as described previously^23^. Thin sections of the same areas imaged in SR-SIM were imaged in a Jeol 1200 transmission electron microscope (JEOL, Japan).

### Plasmodium Assays

The Cb-EmeraldFP sequence was cloned and transfected into *P. Falciparum* 3D7 parasites, and parasites were cultured in RPMI 1640 supplemented with Albumax (Invitrogen). Schizonts were collected on a bed of 70% Percoll as previously described^24^. The schizonts were allowed to egress in a medium containing blood at 1% haematocrit and merozoites at various stages of invasion were fixed in 4% paraformaldehyde with 0.0075% glutaraldehyde as previously described^25^. The junction was stained with a RON4 antibody and the nucleus stained by DAPI. Cb-EmeraldFP labelled actin filaments were fluorescent in the green range. For image acquisition, z–stacks were collected using a UPLSAPO 100× oil (1.40NA) objective on a Delta Vision Core microscope (Image Solutions – Applied Precision, GE) attached to a CoolSNAP HQ2 CCD camera. Deconvolution was performed using SoftWoRx Suite 2.0 (Applied Precision, GE). An Elyra S1 microscope with Super resolution Structured Illumination (SR-SIM) (Zeiss) was used for super-resolution dissection of AMA1 staining on the merozoite surface

P. knowlesi A1-H.1 parasites were cultured as described previously^26^ in human O+ RBCs in RPMI-HEPES media supplemented with 2.3 g/L sodium bicarbonate, 2 g/L dextrose, 0.05 g/L hypoxanthine, 0.025 g/L gentamicin, 0.292 g/L L-glutamine, 5 g/L Albumax II (Gibco) and 10% (v/v) equine serum (Gibco). Parasites were cultured at 37 °C with a gas mixture of 90% N2, 5% O2 and 5% CO2.

### P. knowlesi merozoite imaging

PkRON2*mCherry-HA A1-H.1 merozoites were incubated with fresh erythrocytes, shaken at 1000 rpm, 37 °C for 90 secs and samples fixed by addition of an equal volume of 2x fixative (8% paraformaldehyde/0.015% glutaraldehyde, Sigma-Aldrich). Cells were permeabilised with 0.1% Triton X-100/PBS for 10 mins and blocked with 3% BSA/PBS. Cells were incubated with 1:500 mouse anti-actin^26^ and 1:1000 rat anti-HA (3F10, Roche) followed by 1:1000 anti-mouse Alexa 488 (Invitrogen) and 1:1000 anti-rat Alexa 594 (Invitrogen) secondary antibodies. Cells were smeared on slides and mounted in VectaShield (Vector Laboratories) with 0.1 ng/ml DAPI to label the parasite nucleus. Widefield fluorescent microscopy images were acquired with a Nikon plan apo 100x/1.45 oil immersion lens on a Nikon eclipse Ti microscope and images processed with NIS Elements (Nikon).

## Acknowledgments

We acknowledge the assistance of the 3IP (Institute of Infection, Immunity and Inflammation Imaging Platform) at the University of Glasgow. We also want to thank Dr. Philip Thompson for providing us with the reagent BIPPO and Dr. Maryse Lebrun for anti-RON2 antibodies.

## Author contributions

MD, JB, IT and MM conceived, designed, or planned the study. MD, JP, GP, OL, SD, GSP, JS, LG, and FL-B contributed to the acquisition, analysis, or interpretation of the data. MM drafted the manuscript. All authors critically reviewed or revised the manuscript for intellectual content and approved the final version.

## Data and materials availability

Data is available upon request.

---

**Figure S1.**
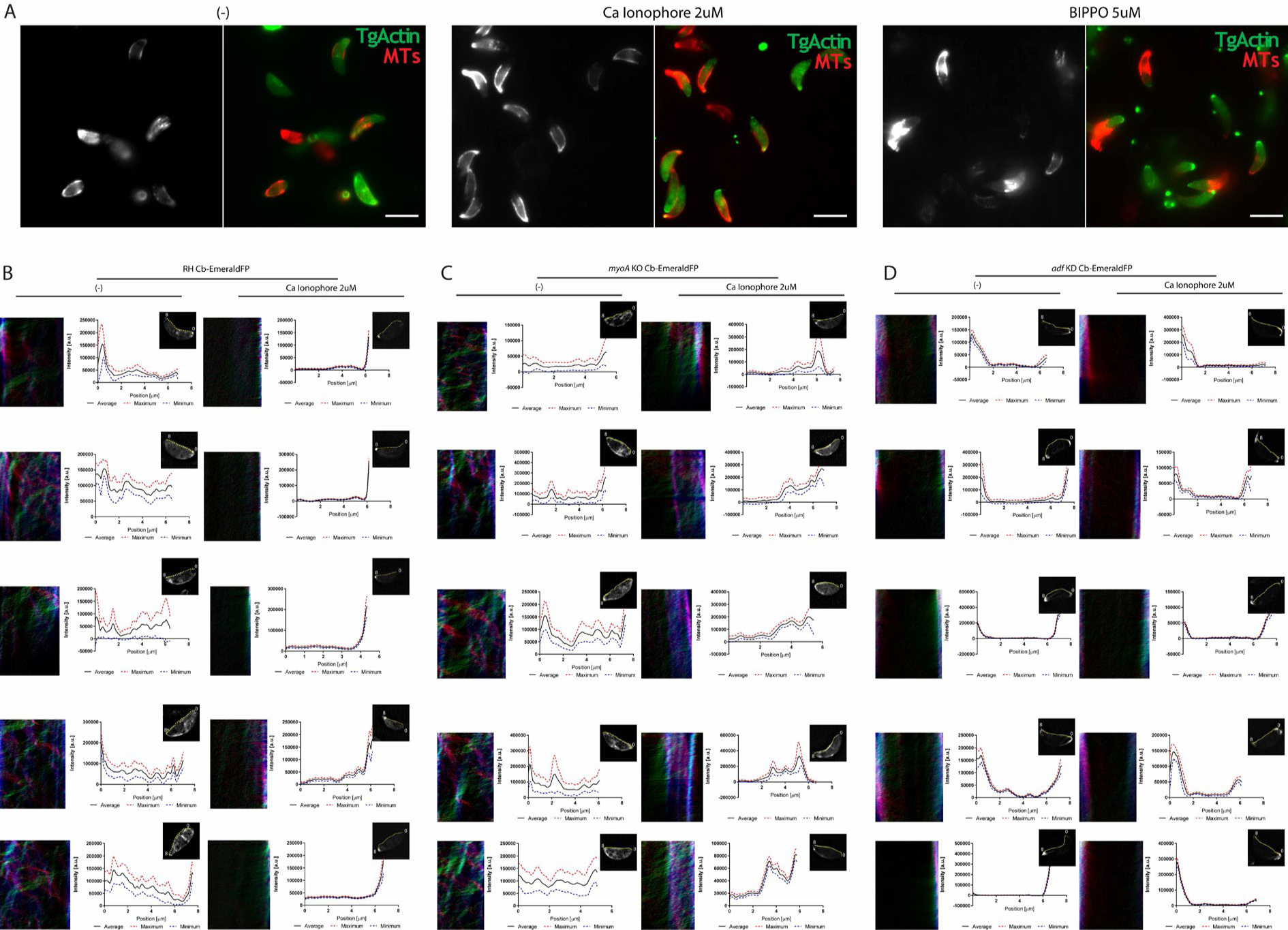
Flow analysis on extracellular parasites and representative images from drug treatment. **A.** Representative images from wild type parasites with calcium ionophore and BIPPO treatment. **B.** Kymograph flow analysis examples done in RH Cb EmeraldFP parasites with and without Ca^2+^ Ionophore treatment. Before treatment, the parasite appears to show dynamic actin behaviour across the entire periphery, yet upon addition of Ca^2+^ Ionophore strong actin accumulation can be observed at the apical tip as represented in the kymograph measurement. **C.** Kymograph flow analysis examples done in *myoA* KO Cb EmeraldFP parasites with and without Ca^2+^ Ionophore treatment. Before treatment, the parasite appears to show dynamic actin behaviour across the entire periphery similar to *wt. U*pon addition of Ca^2+^ Ionophore actin can be observed at the periphery, covering approximately half of the parasite peripherical length as represented in the kymograph measurement. **D.** Kymograph flow analysis examples done in *adf* KD Cb EmeraldFP parasites with and without Ca^2+^ Ionophore treatment. Before treatment, the parasite appears to lack dynamic actin behaviour across the entire periphery unlike *wt* and *myoA* KO, upon addition of Ca^2+^ Ionophore no further change is detected. The colour-coded kymograph represents forward movement (red), backwards movement (green) and static (blue).

**Figure S2.**
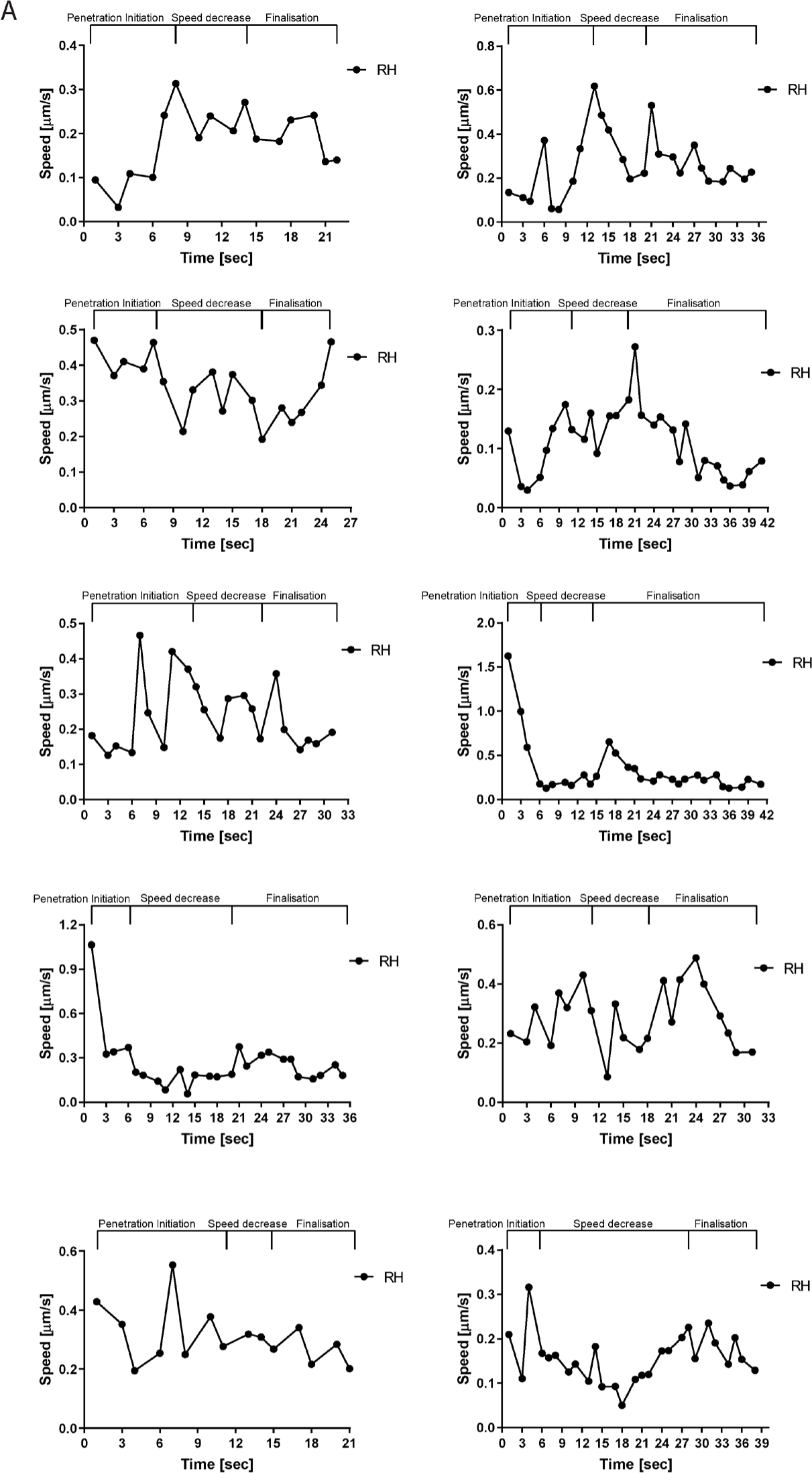
Speed profiles of penetrating parasites. 10 examples of speed profiles of penetrating parasites as shown in Figure 2.

**Figure S3.**
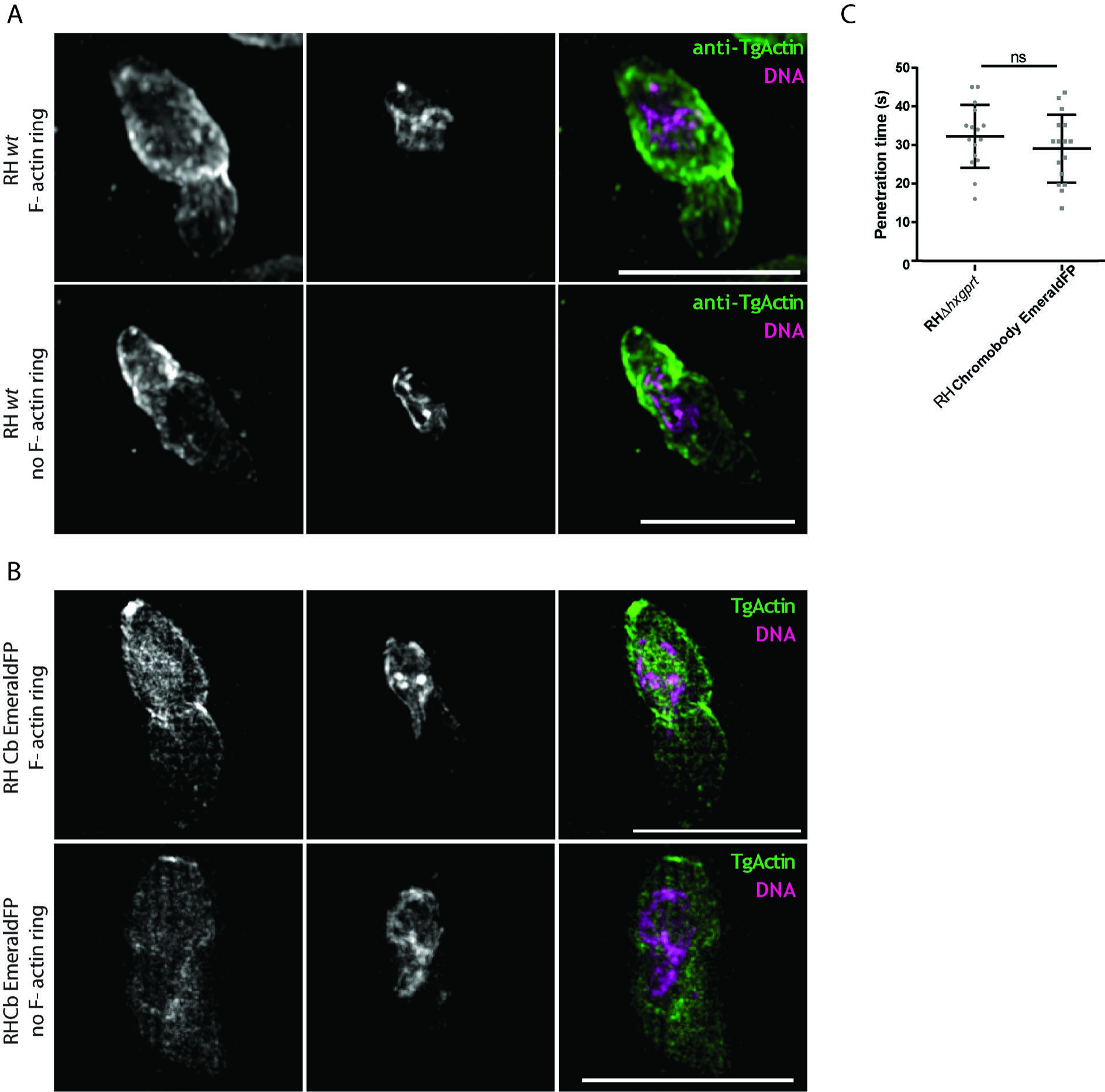
Super resolution microscopy during invasion between wt and chromobody EmeraldFP expressing parasites. **A.** SR images depicting wild type parasites using anti-actin antibodies. In the upper panel a case with an actin ring is shown. In the lower panel, no F-actin ring can be observed. **B.** SR images depicting wild type parasites using chromobodies against actin. In the upper panel a case with an actin ring is shown. In the lower panel, no F-actin ring can be observed. **C.** Comparison of invasion speeds of RH wild type parasites and parasites expressing Cb-EmeraldFP. 15 movies of each strain were analysed in FIJI ImageJ. One-way ANOVA analysis was performed for each graph. P-value <0.0001.

**Figure S4.**
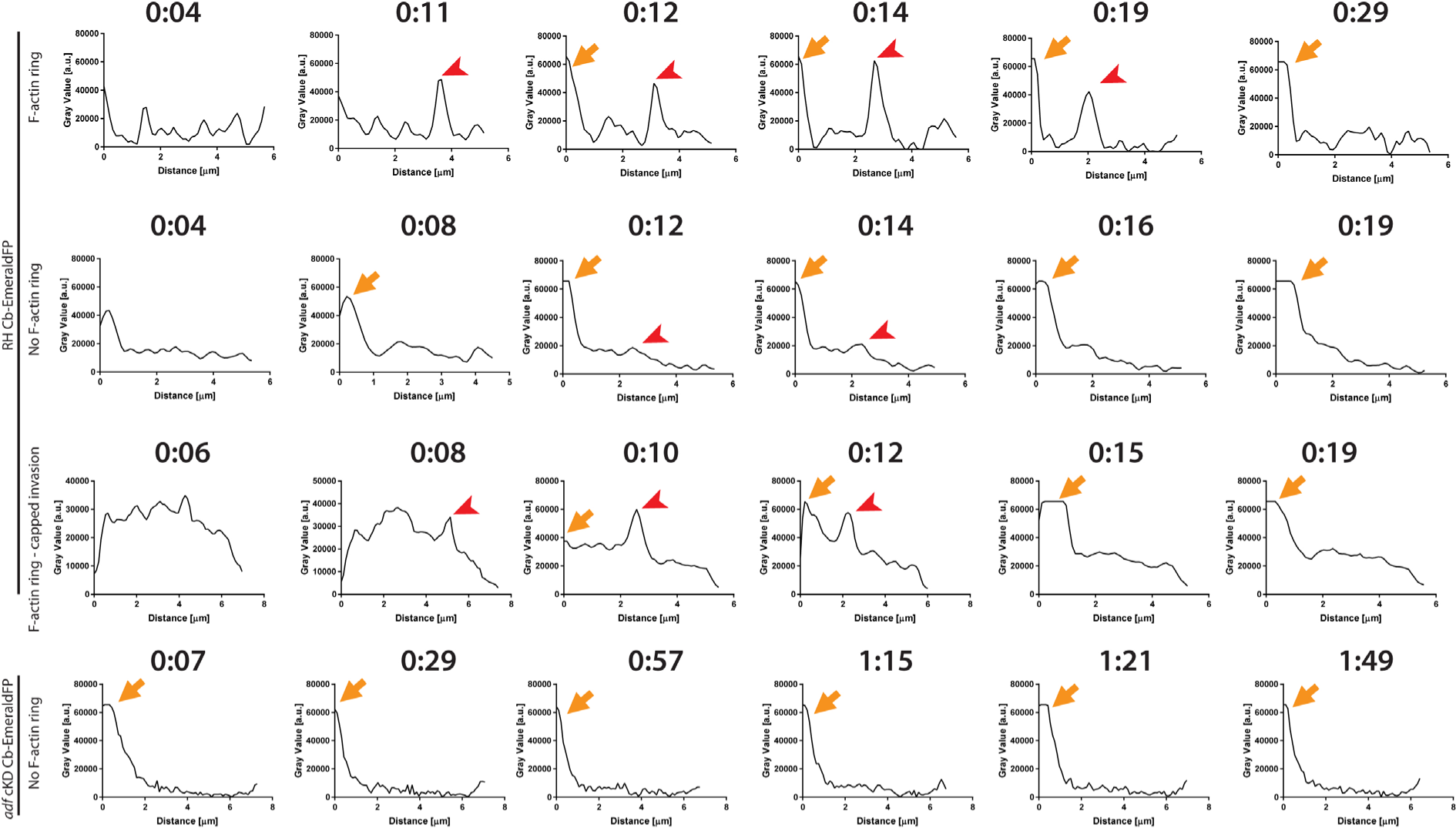
F-actin dynamics during invasion and intensity profiles. Intensity plot profiles on Time-lapse analysis of invading RH Cb-EmeraldFP parasites into HFFs. Analysis performed as in Figure 3. The plot profile for wildtype parasites forming F-actin ring at the junction shows two distinct peaks. Accumulation of F-actin at the posterior pole corresponds to the orange arrowhead, while F-actin at the TJ corresponds to red arrowhead. In case of invasion events without formation of an F-actin ring only posterior accumulation can be detected. An F-actin ring is also still formed on rare events such as capped invasion. During invasion of *adf* cKD parasites (bottom panel) no F-actin ring is formed. Similar to RH parasites F-actin accumulates at the posterior pole (orange arrow).

**Figure S5.**
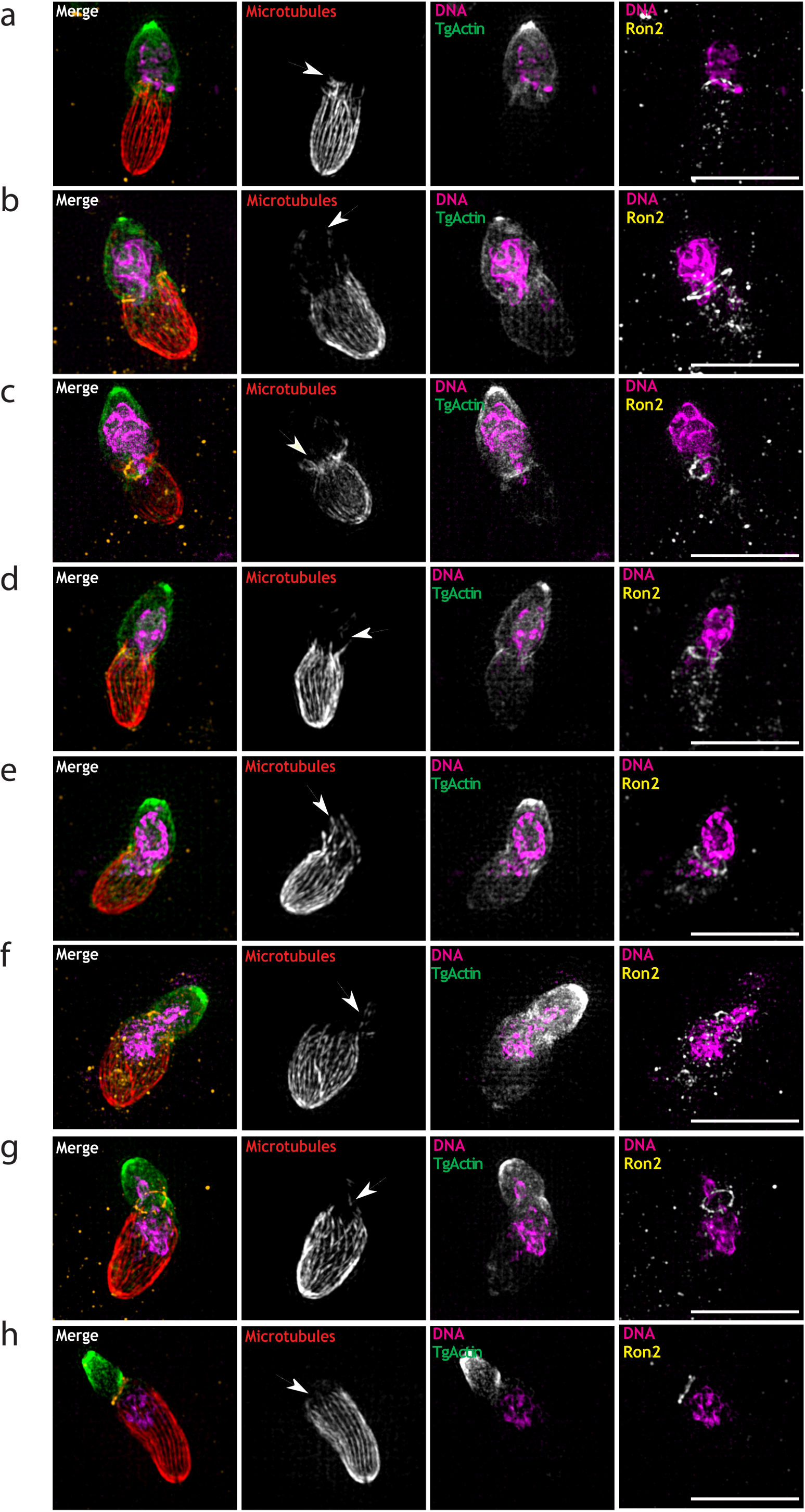
F-actin dynamics and Microtubules during invasion. A-H. SR images of invading parasites in different stages of invasion. The gallery shows invasion in advancing steps. The MTs are deformed during invasion as show across the gallery. In certain cases, such as (b-d), the MTs seem to extend further than usual closely aligned to the nucleus. F-actin forms a mesh that is more prevalent in the portion of the parasite that is still outside. The nucleus is constricted when it is going through the TJ (e-g).

## Supplemental Movies

**Movie S1. SR-SIM time-lapse of actin flow with skeletonisation analysis.** Time-lapse of SR-SIM showing RH, *myoA*KO and *adf*KD parasites expressing chromobody EmeraldFP. Imaging speed 7 fps.

**Movie S2. SR-SIM time-lapse of actin flow before and after calcium ionophore A23187 or BIPPO.** Time-lapse of SR-SIM showing RH, *myoA*KO and *adf*KD parasites expressing chromobody EmeraldFP before and after calcium ionophore A23187 or BIPPO. Imaging speed 7 fps.

**Movie S3. Wild type RH parasites invading HFF cells.** Two time-lapse moving showing live invasion of RH wild type parasites. Imaging speed 7 fps.

**Movie S4. Mutants parasite lines invading HFF cells.** Parasite mutant lines *myoA* KO, *act1* KO and *mlc1* KO time-lapse moving showing live invasion of *myoA* KO parasites. Imaging speed 7 fps.

**Movie S5. Abortive invasion by different parasite strains.** Wild type RH, *myoA* KO, *act1* K and *mlc1* KO parasites time-lapse moving showing live abortive invasion of RH wild type parasites. Imaging speed 7 fps.

**Movie S6. F-actin events during invasion.** An F-actin ring appears during initial parasite penetration. However, the presence of an F-actin ring is not required in some cases. In rare cases such as capped invasion, the F-actin ring is also present. In mutants with abrogated F-actin dynamics such as *adf cKD*, the F-actin ring is not present during invasion. Time-lapse moving showing live invasion of Cb-EmeraldFP parasites. The signal is F-actin Cb-EmeraldFP. Imaging speed 7 fps.

**Movie S7. RH Chromobody EmeraldFP parasites invasion and F-actin dynamics during nucleus passage.** Nucleus passage when the F-actin ring is present and absent. Time-lapse moving showing live invasion of Cb-EmeraldFP wild type parasites with an F-actin ring formation. Cb-EmeraldFP is shown in green, Hoechst staining is shown in magenta. Imaging speed 7 fps.

**Movie S8. 3D rendering of RH and *adf cKD* Chromobody EmeraldFP parasites invasion with and without F-actin ring.** 3D rendering using IMARIS software on invading Cb-EmeraldFP wild type and *adf cKD* parasites with an F-actin ring formation. Cb-EmeraldFP is shown in green, Hoechst staining is shown in magenta. Imaging speed 7 fps.

**Movie S9. 3D rendering of RH Chromobody EmeraldFP parasites invasion with microtubule and TJ staining.** 3D rendering using IMARIS software on invading Cb-EmeraldFP parasites with an F-actin ring formation. Cb-EmeraldFP is shown in green, Hoechst staining is shown in magenta, RON2 staining is show in yellow and MT staining is show in red. Imaging speed 7 fps.

